# Examples of adaptive peak tracking as found in the fossil record

**DOI:** 10.64898/2025.12.07.692820

**Authors:** Rolf Ergon

## Abstract

Species that have persisted over millions of years have done so because they have been able to track peaks in an adaptive landscape well enough to survive and reproduce. Such optima are defined by the mean phenotypic values that maximize mean fitness, and they are predominantly functions of the environment, for example the sea temperature. The mean phenotypic values over time will thus predominantly be determined by the environment over time, and the trait history may be found in the fossil record. Here I use fossil data from four cases found in the literature, and show that adaptive peak tracking models give better results than alternative weighted least squares and directional random walk models. The model performances are compared by use of weighted mean squared errors and Akaike information criterion results.

## 1 Introduction

Adaptive peak tracking is known from various engineering applications, but tracking models are also relevant in evolutionary biology settings. They are then based on an adaptive landscape metaphor, which assumes that the mean population fitness in a population is a function of possibly multivariate mean phenotypic traits, with an optimum (adaptive peak) for a specific set of mean traits. See Pigliucci (2013) for historical background and clarifying discussion. Below, I also make use of an individual fitness landscape spanned by axes for individual phenotypic values, for example body size or number of fin rays. Here, individual fitness is a measure of reproductive success, i.e., to which degree genes are passed on to future generations. For species with sexual reproduction the phenotypic values are midparent values, i.e., the means of male and female traits.

It has for a long time been known that populations and species of larger size are found in colder environments, while populations and species of smaller size are found in warmer regions. This is called Bergman’s rule after the German biologist Carl Bergman, who described the pattern in 1847. Although there are exceptions from this rule, it is today referred to also in cases where a gradual change in temperature leads to gradual changes in body size, and it is then at least implicitly assumed that the body size tracks a peak in an adaptive landscape. As shown by the last real data example in Section 3, such tracking of environmental change is not restricted to body size, although it indirectly may be so.

Adaptive peak tracking in evolutionary biology can be described by the block diagram in Fig. 1. From a possibly multivariate environment *u*(*t*) follows a peak *θ*_*i*_(*t*) in the individual fitness landscape, i.e., individual phenotypic values that maximize fitness, and when *u*(*t*) changes the position of this peak will change accordingly. This will again result in a moving adaptive peak *θ*_*p*_(*t*) in the adaptive landscape, which with a non-symmetrical individual fitness function will have a different position than *θ*_*i*_(*t*). As discussed in Ergon (2025), the two peaks may also have different locations owing to constraints on the phenotypic values, and conceivably also for other reasons. The essential feature of the model in Fig. 1 is that the tracking process seeks to keep the mean phenotypic trait 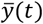 equal to the adaptive peak value *θ*_*p*_(*t*), and from a feedback control point of view it will do so with integral action. In each of the real data cases in Section 3, we will find that *θ*_*p*_(*t*) is a nearly linear function of a dominating environmental driver, which means that also 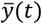 is a linear function of *u*(*t*). As will be seen in these cases, the environmental variables may be quite noisy, and I will therefore use moving average smoothed signals as *u*(*t*). This is also a way to handle errors in the estimated age of the fossils.

**Figure 1.**
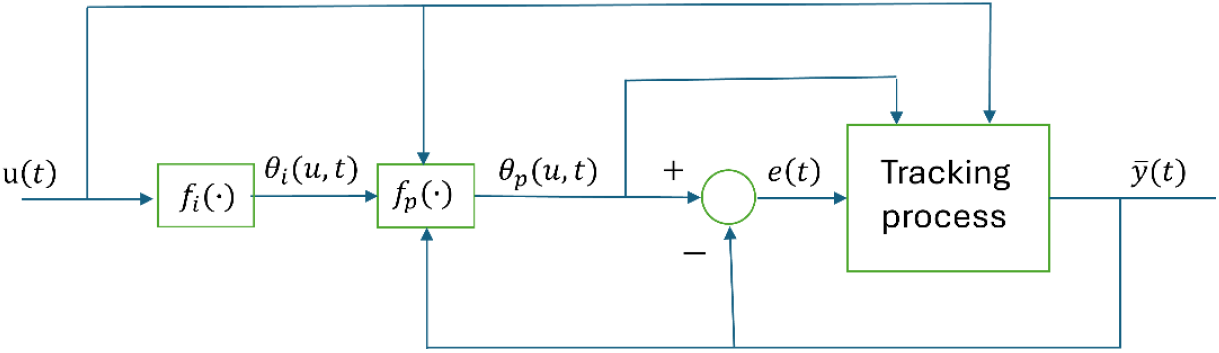
Block diagram for adaptive peak tracking system in evolutionary biology.

The adaptive peak tracking model in Fig. 1 can be applied on all types of evolutionary systems, but in the following I will focus on systems involving fossil data. One example is found in Liow et al. (2024), with fossil data over 2.3 million years on the size of feeding zooids (autozooids or AZ) in encrusting cheilostome bryozoans. The data were collected from 985 fossil colonies as shown in Fig. 2, with a large total number of individual bryozoans. Results in the form of nine *log AZ area mean* values with standard errors were analyzed in Ergon (2025), see panel B in Fig. 3, with present time as zero point. Two samples were omitted for reasons as discussed in Ergon (2025).

**Figure 2.**
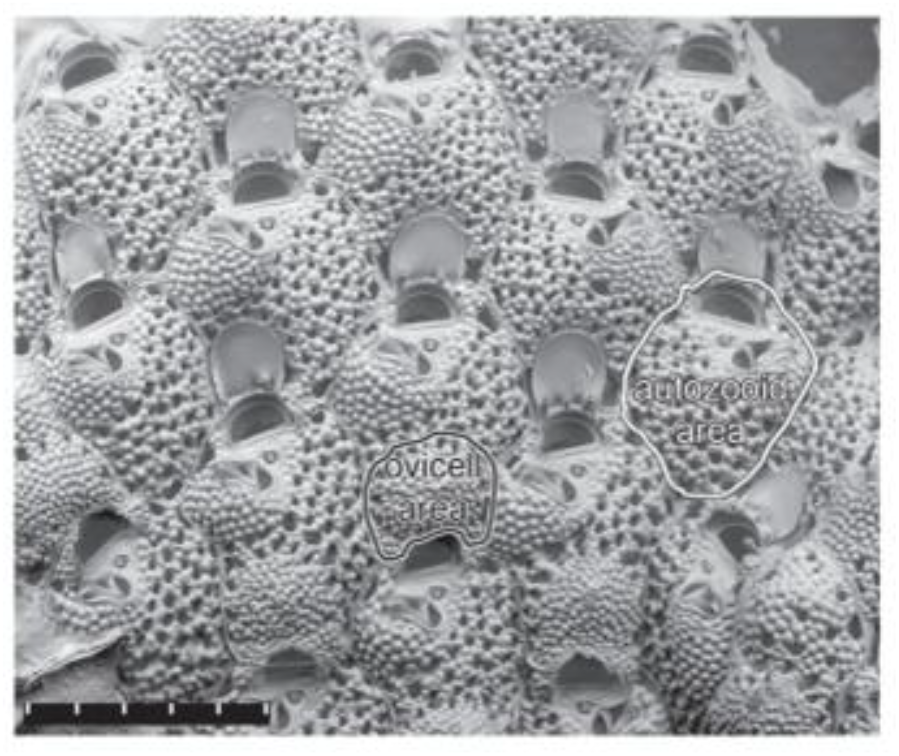
Scanning electron micrograph showing part of a *Microporella agonistes* colony, with perimeters of autozooid (AZ) and ovicell (OV) areas outlined. The length of the scale bar is 500 *μm*. From Liow et al. (2024).

**Figure 3.**
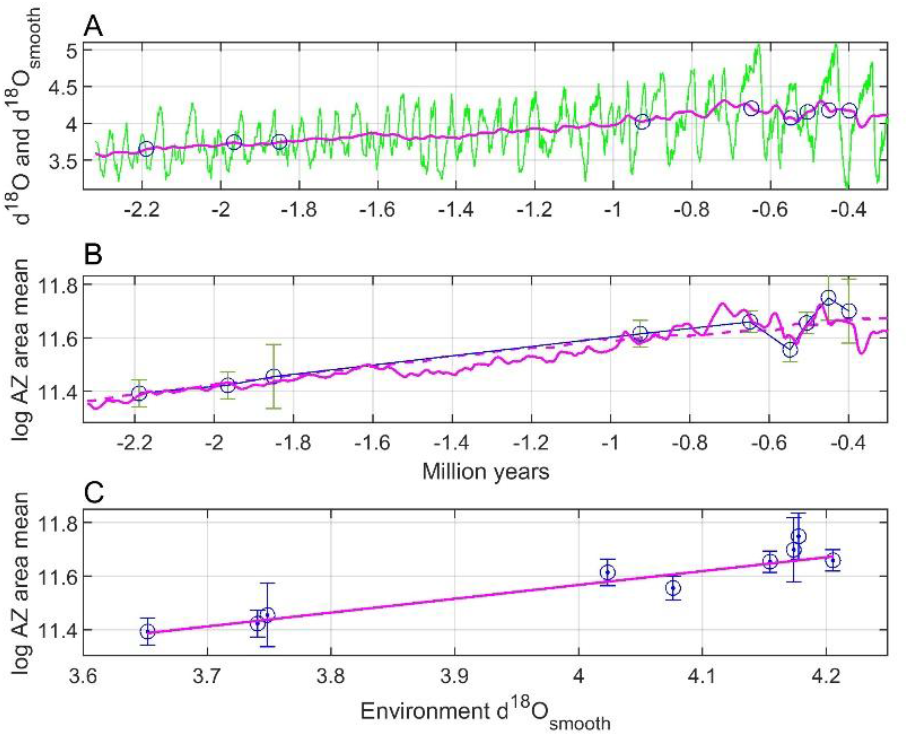
Results for the bryozoan species *Microporella agonistes*, as shown in Fig. 2. Panel A shows raw *∂*^18^*O*(*t*) data (green line) and a centered moving average smoothed version *∂*^18^*O*_*smooth*_(*t*) with window size 100 samples (red line). For the four last samples, the time window size is 0.1 million years, while it is 0.2 million years for samples 4 and 5, and 0.25 million years for samples 1 to 3. Panel B shows *log AZ area mean* with error bars, as found from Fig. S2 in Liow at al. (2024) (circles), as well as predictions (red line). Panel C shows the estimated adaptive peak versus environment function (red line) computed by the weighted least squares method. Panel B also shows predictions with window size 800 samples (dashed red line).

In addition to mean trait standard errors, as shown by error bars in Fig. 3, there are considerable errors in the nine time points, i.e., in the estimated age. This is because mean trait values are found as mean values of mean values for various numbers of colonies found in nine different geological formations with various time spans, and the nine time points are estimated midpoints for these formations. The dominating environmental driver is the sea temperature, with oxygen isotope *∂*^18^*O*(*t*) values used as proxy (Lisiecki and Raymo, 2005). The *∂*^18^*O*(*t*) values increase when the temperature decreases, and they can be converted to temperature by linear transformations (Evans et al., 2024). Because of the potentially large age errors in the mean phenotype data, Ergon (2025) used *u*(*t*) = *∂*^18^*O*_*smooth*_(*t*) as a centered moving average smoothed version of *∂*^18^*O*(*t*). Using the moving average window size as a tuning variable, the nine mean phenotypic values fell quite nicely on a straight line (Fig. 3, panel C), and from weighted least squares (WLS) parameter estimation thus followed continuous predictions of the mean trait values (Fig. 3, panel B). Again, note that a high *∂*^18^*O*(*t*) value means low temperature, and that the AZ area thus in accordance with Bergman’s rule increases when the temperature decreases. See Section 3 for detailed results regarding the ovicell (OV) trait as shown in Fig. 2.

The adaptive peak tracking model in Fig. 1 is useful only if the following error *e*(*t*) is kept sufficiently small, such that the mean phenotypic value stays in the vicinity of the adaptive peak. As referred to in Ergon (2025), there is a lot of support for this assumption. A persistent system excitation in the form of short-term environmental fluctuations as exemplified in Fig. 3, panel A, makes it unlikely that a tracking system gets stuck in a local optimum or in a valley in the adaptive landscape. As illustrated in simulations in Ergon (2025), it is more likely that the mean phenotypic values fluctuate around global optima.

Under the assumption that the following error *e*(*t*) in Fig. 1 is kept sufficiently small, such that 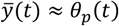, the adaptive peak function *θ*_*p*_(*t*) = *f*(*u*(*t*)) can be identified from *u*(*t*) and 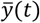 data, and that is especially simple when this function is linear. As illustrated in Fig. 3 it is then possible to find the adaptive peak function from few and irregularly spaced samples, and thus also to predict the mean trait values 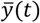. Note that this can be done without detailed knowledge of the tracking process in Fig. 1.

Section 2 describes the methods used in the article. As also used in Liow et al. (2024), a well-established method for analyses of fossil time series data is to find optimal biased or unbiased random walk models (Hunt, 2006). Biased or general random walk (GRW) models are also known as directional evolution models (Hunt, 2012). An alternative model for directional evolution is to use WLS directly on the 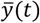 data. In the real data cases in Section 3 I will thus compare three models, (i) tracking with use of WLS on the adaptive peak function 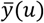, (ii) WLS directly on 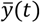, and (iii) GRW. The performances of these models will be compared by use of weighted mean squared errors (WMSE). The models are also compared by use of the Akaike Information Criterion (AIC), which is often used for comparison of different random walk models of fossil data (Hunt, 2006).

In Section 3, the different methods are applied on four real case data sets in the fossil record. The first case is from the already introduced field study of the bryozoan species *Microporella agonistes*, with results for *log AZ area mean* as shown in Fig. 3. The second case is from the same bryozoan study, but with results for *log OV area mean*. The third case is a study of the ostracod species *Poseidonamics major* in Hunt and Roy (2006), while the fourth is a stickleback fish study in Bell et al. (1985), which shows that tracking of environmental change is not restricted to body size traits (although it indirectly may be so). Finally follows a summary and discussion in Section 4.

## 2 Theory and Methods

### 2.1 Essential elements of the tracking process

As discussed in the introduction, it is not necessary to know the details of the tracking process in Fig. 1, but it must have two essential elements. First, there must be a mean fitness function such that 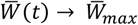 when 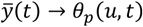. This function defines the adaptive peak, and in the univariate case it can for example be (as in Ergon, 2025)

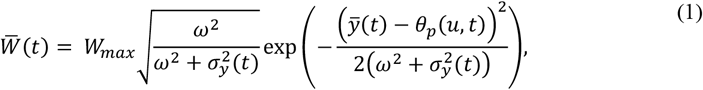

where 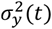 is the variance of *y*(*t*), while *ω*^2^ is the width of the underlying individual fitness function. Second, there must be a selection mechanism, such that individuals with high fitness get the best chance for reproduction (Lande, 1979). A simple example of such a set of selection equations is given in Ergon (2025). This will in most cases result in slow evolution, but owing to plasticity there may still be rapid changes in the mean trait 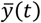 (Ergon, 2025). In addition, mutations may give sudden and positive jumps in the individual fitness (Ch. 7, Walsh and Lynch, 2018).

### 2.1 Weighted least squares estimation for tracking models

In the tracking method, least squares predictions are found as 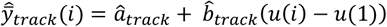, where *u*(*i*) is a moving average smoothed version of the *∂*^18^*O*(*i*) measure at the *N* sampling points. Here, the parameters are found as

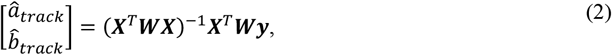

where 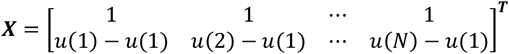 and 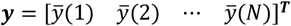, while ***W*** is a diagonal matrix with weights 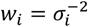 that are the reciprocals of the measurement variances. From this follows continuous predictions

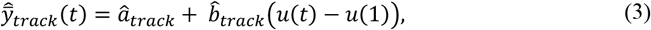

such that 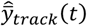 becomes a smoothed and scaled version of the environmental variable.

### 2.2 Weighted least squares estimation for direct WLS models

In the direct WLS method, least squares predictions are found as 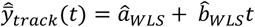. Here, the parameters are found as in Eq. (2), except that 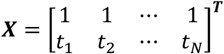, where *t*_*i*_ are the sampling time points.

### 2.3 Random walk models

Hunt (2006) developed what he called a general random walk (GRW) model for analyses of fossil time series, later referred to as a model for directional evolution (Hunt, 2012). This model assumes a random walk process, where at each timestep an increment of evolutionary change in a mean trait value is drawn at random from a distribution of evolutionary steps. It is also assumed that the incremental change is normally distributed with mean value *μ*_*step*_ and variance 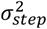. For an observed evolutionary mean trait change Δ*Y* over T time steps, the log-likelihood function for a GRW process is given by (Hunt, 2006)

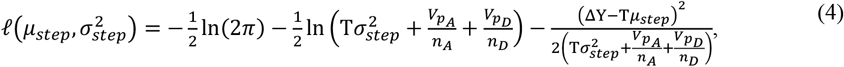

where *n*_*A*_ and *n*_*D*_ are the numbers of observed ancestors and descendants, respectively, while 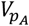 and 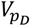 are the corresponding population phenotypic variances. With *N* irregular and sparse samples of mean trait values, multiple ancestor-descendant trait differences over *N* − 1 evolutionary transitions may be used jointly to estimate *μ*_*step*_ and 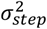 by summing the log-likelihoods according to Eq. (4) over the transitions, i.e., by use of 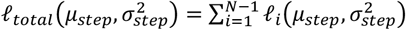. From this expression, step parameters can be found by means of maximum likelihood estimation, and the estimated step size 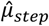 will then be a measure of directional change over time. In the real data examples in Section 3, the optimal GRW parameters are found by use of the MATLAB function *fmincon*, with initial parameter values sat to zero.

The GRW model has an obvious weakness in that it is difficult to estimate 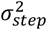 for small values of *T* and large values of 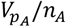 and 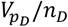. In the three examples in Section 3 the result is 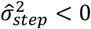, such that the step variance must be set to 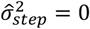.

Eq. (4) is not intended for predictions, but once 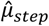 is determined with 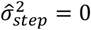, a prediction model

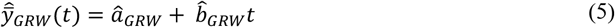

is found by use of 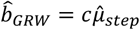, and by fitting a parameter *a*_*GWR*_ to the data by a least squares approach. Here, *c* is a scaling constant that I consistently will set to *c* = 1.

### 2.4 Weighted mean squared errors

Comparisons of responses 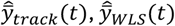 and 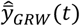 can be done by use of weighted mean squared errors

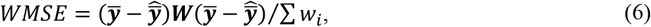

where 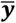 and 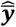 are *N* × 1 data vectors, and where ***W*** is a diagonal matrix with the weights *w*_*i*_. Note that we in the special ordinary least squares (OLS) case have ***W*** = ***I*** and ∑ *w*_*i*_ = *N*. I thus assume independent and normally distributed random variables 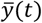. Note that when the weights are the reciprocals of the measurement variances, as done here, the prediction slope parameters 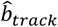 and 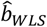 are the best linear unbiased estimates (BLUE).

### 2.4 Theory for the Akaike Information Criterion

Model performances may be compared by use of the Akaike information criterion (AIC), and the model with the lowest AIC value should then be chosen (Hunt, 2006). For GRW, the AIC value with compensation for short data is given by (Hunt, 2006),

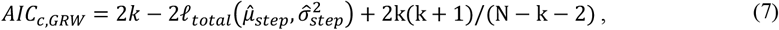

where k = 2 is the number of estimated parameters, while *N* is the number of samples (the number of transitions is N − 1). Since in all the real data examples in Section 3 the estimated variance 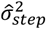 is negative, the realistic value of *AIC*_*c,GRW*_ is found by finding 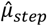 with 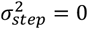.

For comparisons with tracking and WLS models, Eq. (7) must be replaced by (Banks and Joyner, 2017)

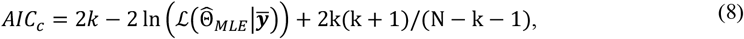

where 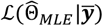 is the likelihood of the estimated parameters using maximum likelihood estimation, given the vector 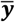 of observed data. Banks and Joyner (2017) further assume a statistical model *Y*_*i*_ = *f*(*t*_*i*_, ***q***_0_) + *ω*_*i*_ℰ_*i*_, where ***q***_0_ is a parameter vector, while *ω*_*i*_ are known weights. Here, *ε*_*i*_ for *i* = 1, 2, …, *N* are i.i.d. 𝒩(0, *σ*^2^), such that *σ*^2^ must be estimated together with ***q***_0_. In the present setting the weights are 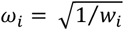, and the final *AIC* expression in Banks and Joyner (2017) thus becomes

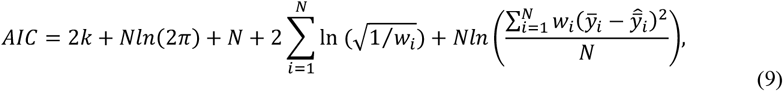

from which *AIC*_*c,track*_ and *AIC*_*c,WLS*_ follow by adding 2k(k + 1)/(N − k − 1). Here, *k* = 3, since also the underlying parameter *σ*^2^ must be estimated in addition to *a*_*WLS*_ and *b*_*WLS*_. Note that 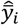 in Eq. (9) is the result of either a tracking model or WLS, and that Eq. (9) therefore is not valid with use of 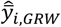 as found from Eq. (5).

Two models with known AIC values can be compared by use of Akaike weights (Wagenmakers and Farrell, 2004). Assuming that that the tracking model is best, as we will find in the real data examples in Section 3, we first compute

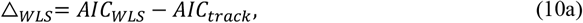

and

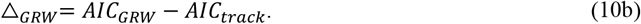

We then find the Akaike weights

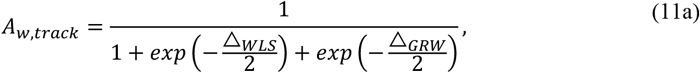

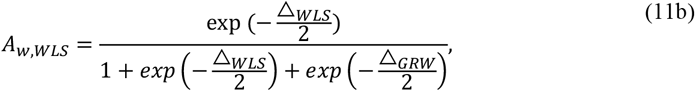

and

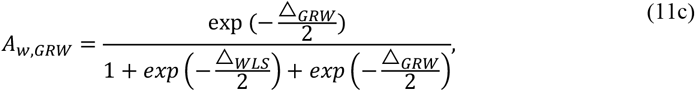

The sum of these weights is one, and the weights can be directly interpreted as conditional probabilities for the models (Wagenmakers and Farrell, 2004).

## 3 Real data cases

### 3.1 Bryozoan AZ case

As already introduced, Liow et al. (2024) presented results from a field study of the bryozoan species *Microporella agonistes*, based on a fossil record over 2.3 million years. The scanning electron micrograph in Fig. 2 shows perimeters of autozooid (AZ) and ovicell (OV) areas, and Fig. 3 shows predictions of *log AZ area mean* as found in Ergon (2025). I used the environmental driver *∂*^18^*O*(*t*) as found in Lisiecki and Raymo (2005), and the optimal moving average window size was 100. In these data the time resolution for *t* > −0.6 million years is 0.001, such that the moving time window size is 0.1 million years. For −1.5 < *t* < −0.6 the time window size is 0.2 million years, while it for *t* < −1.5 is 0.25 million years. With 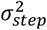 as a free variable, the term 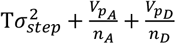 in Eq. (4) for one or several of the transitions becomes negative in the course of the parameter search, such that the log-likelihood becomes a complex number. The search is therefore limited to 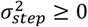, resulting in 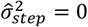. See Table A1 in Appendix A for data, and Fig. 4 and Table 1 for results. Note that present time is used as zero point. Also note that there are some unfortunate errors in the AIC results given in Ergon (2025).

**Table 1.** Prediction slope, WMSE and AIC results for Bryozoan AZ case.

|  | Tracking model | WLS model | GRW model |
| --- | --- | --- | --- |
| Prediction slope | — | 0.1519 | 0.1441 |
| WMSE | 0.0010 | 0.0019 | 0.0019 |
| $AIC_c$ | −23 | −18 | −10 |
| Akaike weights | 0.9372 | 0.0613 | 0.0015 |

**Figure 4.**
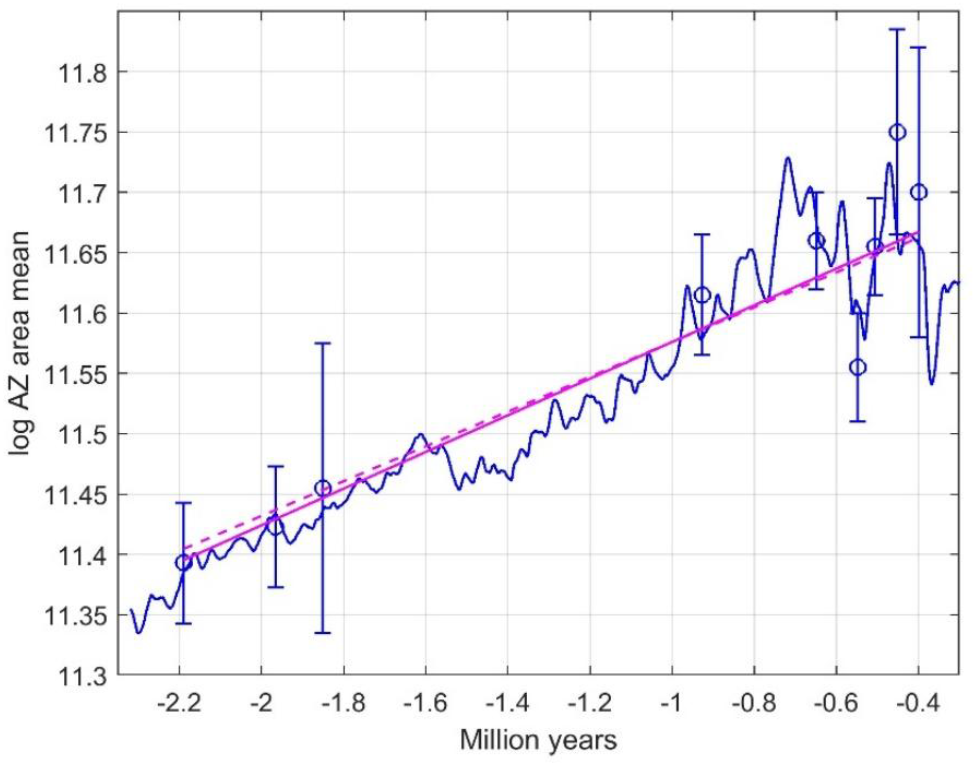
Prediction results for Bryozoan AZ case, with mean trait observations (circles and error bars), tracking predictions (solid blue line), WLS predictions (red line), and GRW predictions with 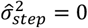 (dashed red line).

### 3.2 Bryozoan OV case

Fig. 5 shows results corresponding to Fig. 4, but now for *log OV area mean*. The optimal moving window size was here 700 (0.7 to 1.75 million years). The parameter search was also in this case limited to 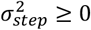, resulting in 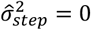. See Table A2 in Appendix A for data, and Fig. 5 and Table 2 for results.

**Table 2.**
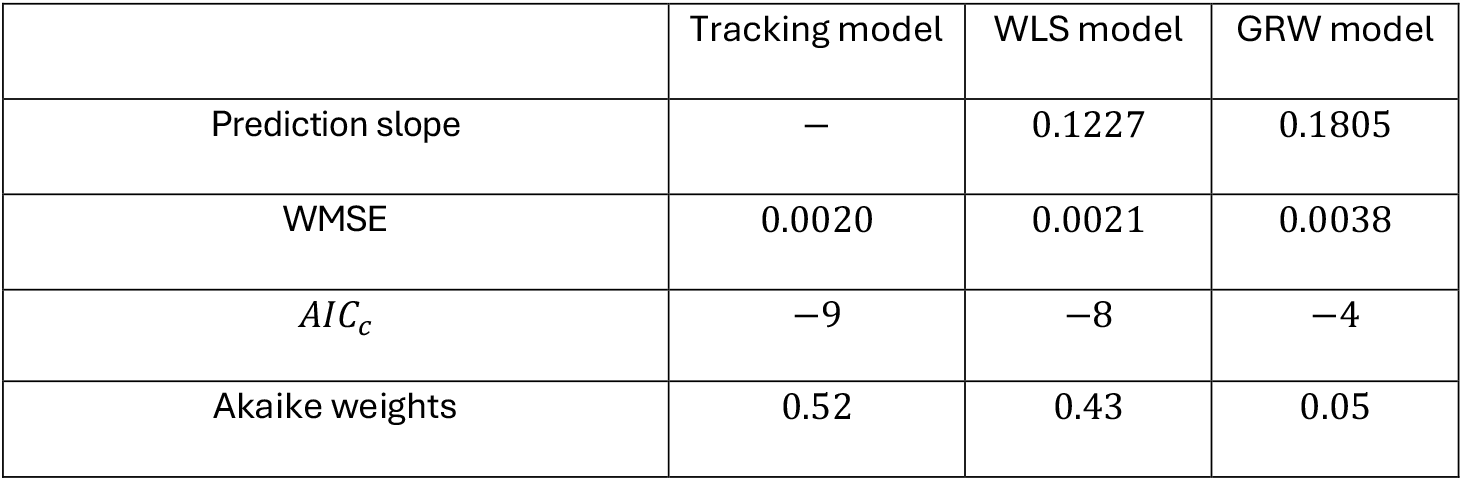
Prediction slope, WMSE and AIC results for Bryozoan OV case.

**Figure 5.**
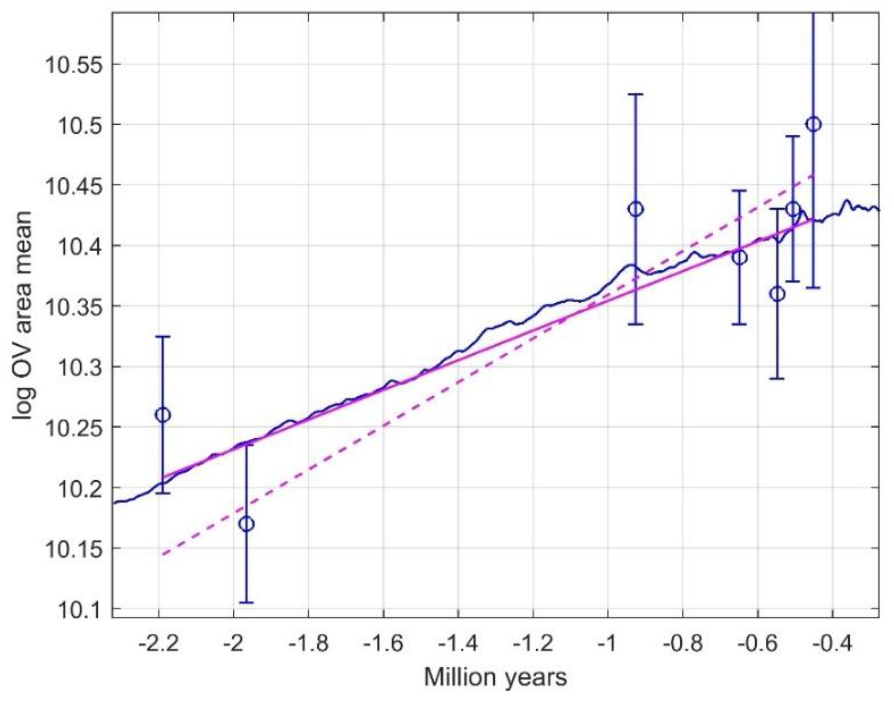
Prediction results for Bryozoan case 1, with mean trait observations (circles with standard error bars), tracking predictions (solid blue line), WLS predictions (red line), and GRW predictions with 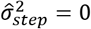 (dashed red line).

### 3.3 Ostracod case

Body size data for the deep-sea ostracod (seed shrimp) species *Poseidonamicus major* are found in Hunt and Roy (2006), Table 4, see Table A3 in Appendix A for details. The body size varies from *y* = 707 to *y* = 876 *μm*, with the variance in the measurements estimated to be 467/*n* (*μm*)^2^, where *n* is the number of samples. The standard error is thus 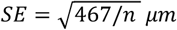, which on a log scale translates to 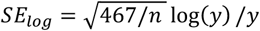.

The data in Hunt and Roy (2006) include Mg/Ca temperatures, but a tracking model based on this information is not any better than the WLS model. Instead, I used the environmental driver *∂*^18^*O*(*t*) as found in Lisiecki and Raymo (2005), and the optimal moving average window size was here 80 samples (0.08 to 0.2 million years). The parameter search was also in this case limited to 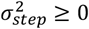, resulting in 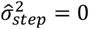. See Table A3 in Appendix A for data, and Fig. 6 and Table 3 for results.

**Table 3.**
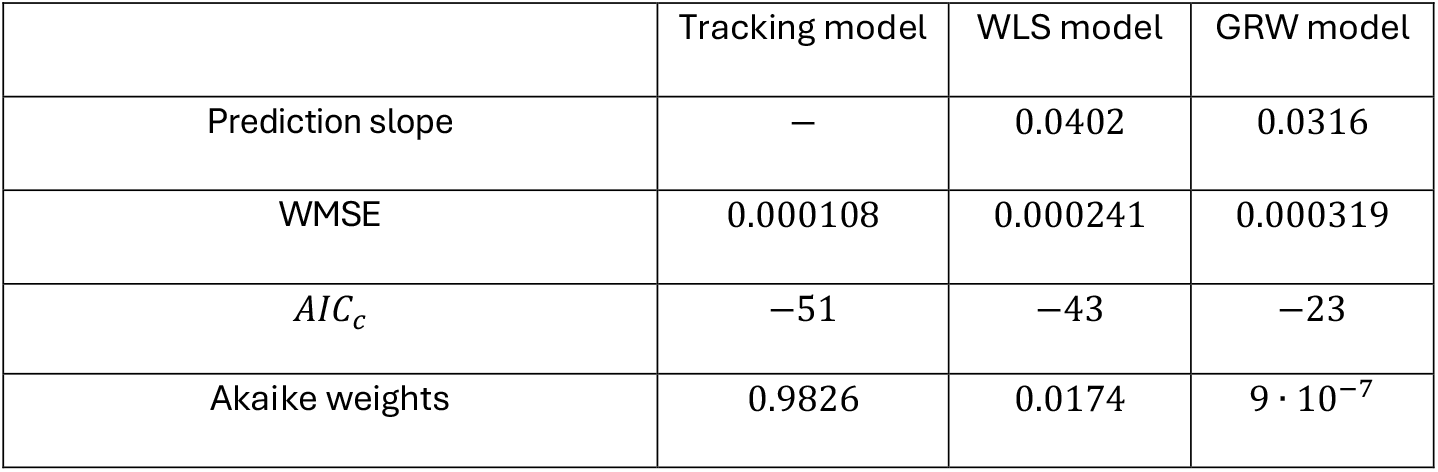
Prediction slope, WMSE and AIC results for Ostracod case.

**Table 4.** Prediction slope, WMSE and AIC results for Stickleback fish case.

|  | Tracking model 1 | Tracking model 2 | WLS model | GRW model |
| --- | --- | --- | --- | --- |
| Prediction slope | — | — | 1.2605 | 1.2531 |
| Zero time point | −9.994 | -10.034 | — | — |
| Time window size | 192 | 196 | — | — |
| WMSE | 0.000216 | 0.000259 | 0.000401 | 0.000401 |
| $AIC_c$ | −131 | −126 | −121 | −104 |
| Akaike weights | 0.9941 | — | 0.0059 | $1.8 \cdot 10^{-6}$ |
| Akaike weights | — | 0.9405 | 0.0595 | $18 \cdot 10^{-6}$ |
| Akaike weights | 0.91 | 0.09 | — | — |

**Figure 6.**
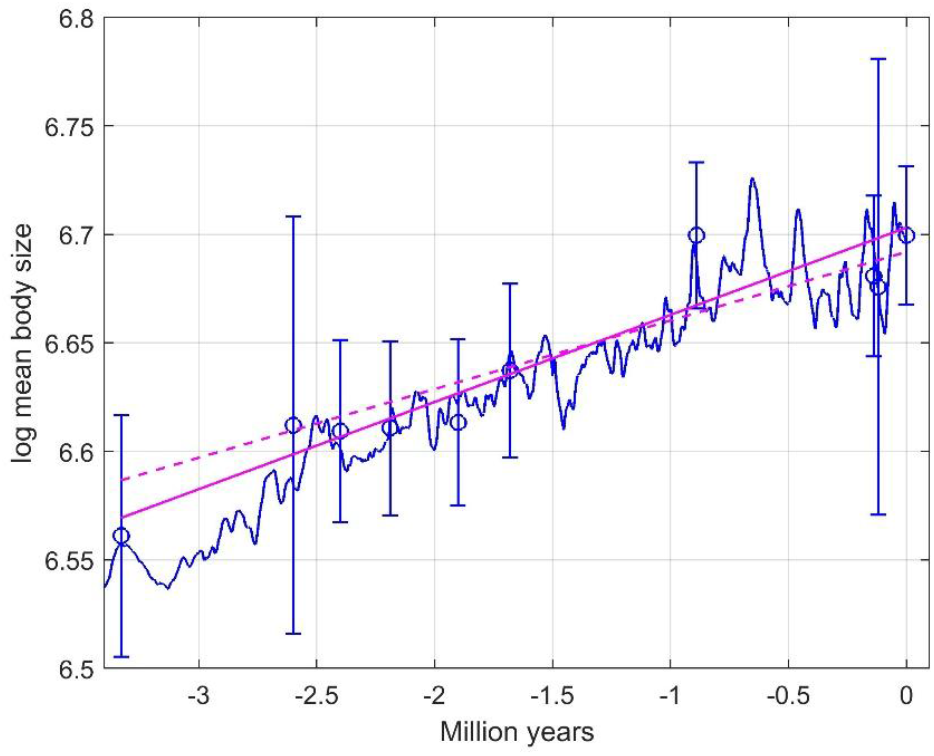
Prediction results for Ostracod case, with mean trait observations (circles with standard error bars), tracking predictions (solid blue line), WLS predictions (red line), and GRW predictions with 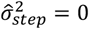 (dashed red line).

### 3.4 Stickleback fish case

In this case I use data that show how the dorsal fin ray number evolved in *Gasterosteus doryssus*, an extinct species of a small freshwater stickleback fish that inhabited inland freshwater habitats of the North American Great Basin during the Miocene (Bell et al., 1985). See Table A4 in Appendix A for data. In the data analysis the mean trait values are log-transformed, with the measurement errors transformed accordingly, i.e., 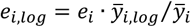, where *e*_*i*_ is found from the given standard deviations *SD* as 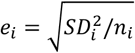.

The original and log-transformed samples thus have equal coefficients of variation.

For a tracking model, the available environmental data was *∂*^18^*O*(*t*) as found in Westerhold et al. (2020), Table S34, column marked ISOBENbinned_d18Ointerp. This is obviously not the direct driver of evolution in what is now Nevada, but there is an association between deep sea temperatures and local Nevada temperatures.

The zero time point not given in Bell et al. (1985), but since *t* = 0 in the data is approximately 10 million years ago (Ma) (personal information) it was possible to align the data in two different ways such that the prediction results had local optima, and we thus have two stickleback fish cases. In both cases optimal moving average window sizes of *n* samples were also found by a manual search using raw data from 9.8 to 10.3 Ma, and since the time resolution is 0.002 million years, the moving time window sizes were *n* · 0.002 million years. In addition to the underlying parameter *σ*^2^ and the elevation and slope parameters there are thus two additional unknown parameters, such that the number of estimated parameters is 5.

- Case 1: The best tracking result was obtained when *t* = 0 in the data corresponds to *t*_0_ = −9.994 million years with present time as zero point (see Figures 7 and 8, and Table 4). Optimal moving average time window size was 0.192 million years.
- Case 2: The zero point *t*_0_ = −10.034 gave almost the same good tracking result (see Figure 9, and Table 4). Optimal moving average time window size was 0.198 million years.

**Figure 7.**
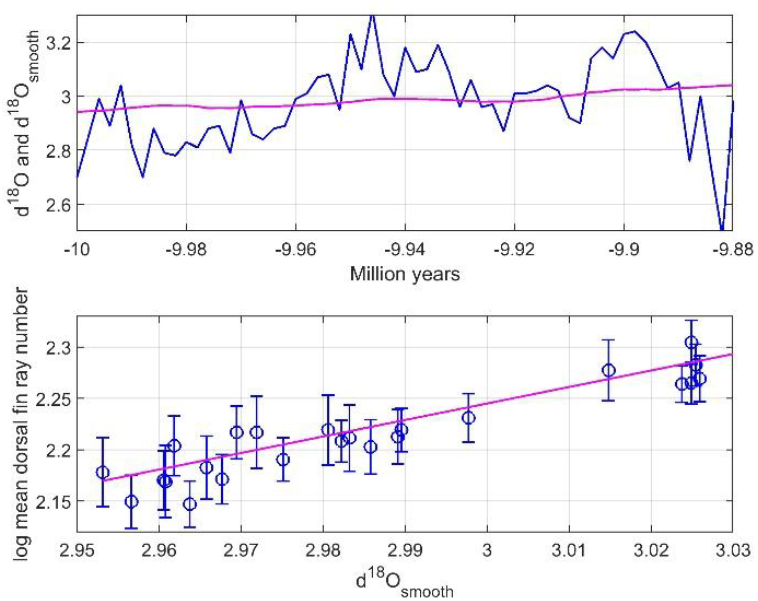
Plots for the Stickleback fish case 1, with upper panel showing *∂*^18^*O*(*t*) (blue line) and *u*(*t*) = *∂*^18^*O*_*smooth*_(*t*) with window size 96 (red line), while the lower panel shows data points 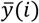 and *u*(*i*) = *∂*^18^*O*_*smooth*_(*i*) with standard error bars, and the estimated linear adaptive peak function (red line).

**Figure 8.**
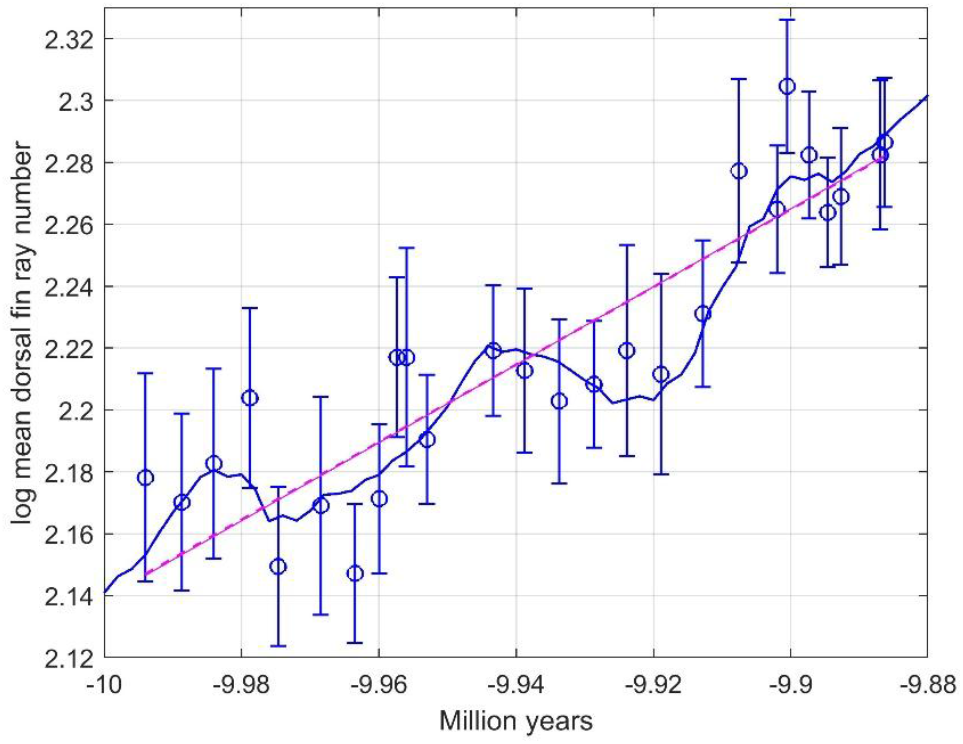
Prediction results for Stickleback fish case 1, with mean trait observations (circles with standard error bars), tracking predictions (blue line), WLS predictions (red line), and GRW predictions with 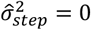 (dashed red line, almost identical to the solid red line).

**Figure 9.**
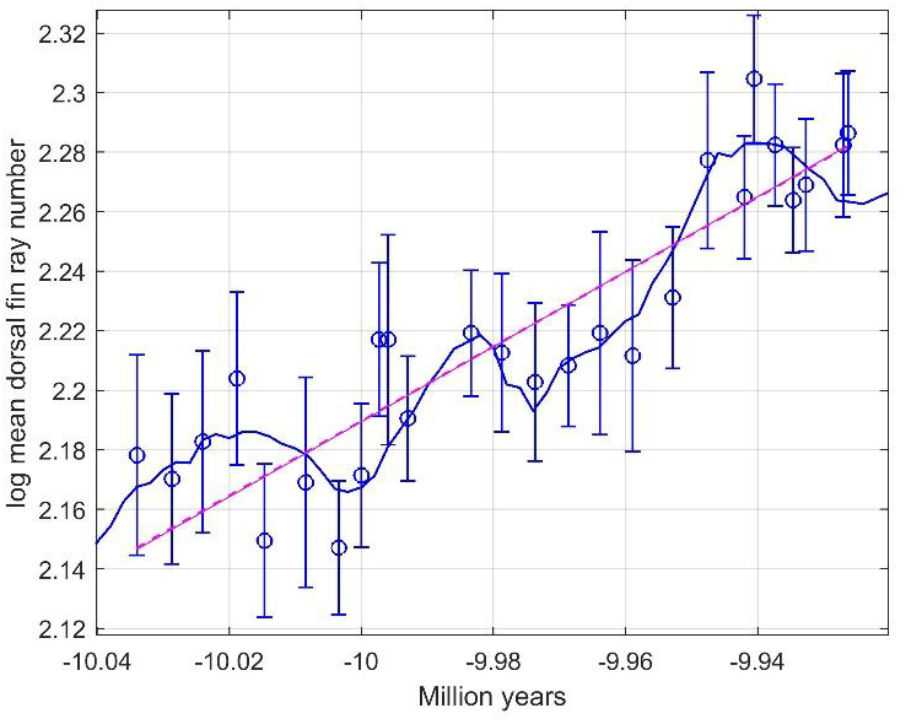
Prediction results for Stickleback fish case 2, with mean trait observations (circles with standard error bars), tracking predictions (blue line), WLS predictions (red line), and GRW predictions with 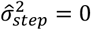 (dashed red line, almost identical to the solid red line).

The time windows are in both cases very wide compared to the time span of 0.1 million years for the data, such that the effect of smoothing mainly is feature extraction, see Section 4 for discussion and Appendix B for details. The parameter search was also in these cases limited to 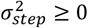, resulting in 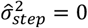.

## 4 Summary and discussion

In this article I use fossil data from four real data cases. I show that in cases with directional evolution, adaptive peak tracking models give better results than alternative weighted least squares (WLS) and general random walk (GRW) models. The model performances are compared by use of weighted mean squared errors (WMSE) and Akaike information criterion (AIC) results.

In all the studied cases, the evolution is driven by a dominating environmental component, but the tracking model in Fig. 1 is in general multivariate. In multivariate cases, the influences from several environmental factors on the adaptive peak movements must be found by causal modeling, which is far beyond the scope of this paper. The adaptive peak may also be influenced by non-symmetrical fitness functions and developmental constraints as discussed in Ergon (2025). In the four real data examples, the positions of the adaptive peaks are approximately linear functions of temperature.

The methodological background is given in Section 2, including the theory for GRW models, and a description of how AIC values are found from WLS based WMSE results. The results for the various models are summarized in Figures 4, 5, 6 and 8, and Tables 1, 2, 3 and 4. Note that the *AIC*_*c,GRW*_ values do not take the estimation of *a*_*GRW*_ in Eq. (5) into account.

In the tracking models I use oxygen isotope data for the environmental drivers. These data were found in deep sea drilling projects, and they can be converted to global ground temperatures through linear transformations (Evans et al., 2024). It is especially interesting that the last case is a study of fossils of a freshwater fish, which verifies that deep sea oxygen isotope data contains information on global temperature. This is perhaps the most convincing case when it comes to tracking model performance (Fig. 8), but it also raises questions as discussed below.

The reason for use of moving average smoothing in the two Bryozoan cases is that the individual data are collected from geological formations with time spans from 0.019 to 0.203 million years, and that there is a lot of variation in the raw *∂*^18^*O*(*t*) data within each time window (Ergon, 2025). As seen in Fig. 3, panel B, smoothing is also a feature extraction method (Herff and Krusienski, 2018), in that the larger window size of 800 samples only captures the overall trend, while the shorter and optimal window size 100 samples reveals more details. Such feature extraction is the main purpose in the Stickleback fish case, where possible errors in the age data are much smaller than in the Bryozoan cases (see Appendix B for details). Note, however, that the raw *∂*^18^*O*(*t*) data as seen in Fig. 7, upper panel, are found from deep-water drilling samples, and that there may be a complex relationship involving various delays, feedback mechanisms and oscillations between these data and the local environment at the field site in what is now Nevada. It is far beyond the aim of this article to consider such complex relationships, and it is also unclear exactly how the local environment affects mean traits. I therefore simply assume that the smoothed deep-sea *∂*^18^*O*(*t*) data captures an essential part of the environmental influence on the mean trait, and that an optimal performance can be found by use of the moving window size and time scale zero point as tuning parameters. This assumption is supported by the prediction results in Figures 8 and 9, and the good AIC results in Table 4. It is also supported by the observation that *∂*^18^*O*_*smooth*_(*t*) appears to be cyclic with an approximate period of 41,000 years, which can be explained by variations in obliquity, i.e., in the Earth’s axial tilt. The relationship between the local obliquity cycles, as experienced by the stickleback fish, and the cycles in the deep-sea drilling samples may as pointed out above be complex, but the essential points are that these cycles have equal frequency but different amplitude and phase, and that these differences are accounted for by the tuning and regression procedures as described.

As shown in Appendix B, replacement of the *∂*^18^*O*_*smooth*_(*t*) input signal by a non-linear function with a sinus term, may in the stickleback fish case give a reduction in WMSE value. However, owing to the increased number of parameters the AIC result will be poorer than for the tracking model. This result supports the evidence that temperature drove the change in dorsal fin ray number.

One may ask whether the obliquity cycles as seen in Figures 8 and 9 are real, or artifacts caused by random noise. The reality of the cycles is seen in the results in Appendix B, Fig. B1, where it, from smoothed and detrended data from 9.2 to 10.8 Ma, is easy to find the period 41,000 years from the total time for 11 cycles. It turns out that that it is more difficult to find the period in a periodogram based on spectral analysis (Fig. B4). The reality of the cycles is also supported by the results in Tian et al. (2013), where obliquity and long eccentricity pacing of the Middle Miocene climate transition is studied.

The stickleback fish case also illustrates an inherent difficulty with the directional evolution concept. The trait evolution as shown in Fig. 8 has an overall positive directional trend, but in several shorter time periods the trend is negative. If the fossils had been 200,000 years younger, it follows from Fig. B1 that the trait evolution instead would have had an overall negative trend, but that the effect of the obliquity cycles would have been the same. This shows that directional evolution is not a system property, but a property of the environment. The essential system property is thus tracking, as illustrated in Fig. 1. In the linear mean trait versus environment cases studied here, the mean trait tracking predictions are simply smoothed and scaled versions of the dominating environmental variable.

In the Bryozoan AZ case, the tracking model was clearly best, while the WLS and GRW models gave very similar results (Fig. 4 and Table 1). The Bryozoan OV case is special in the sense that tracking and WLS models give quite similar results (Fig. 5 and Table 2). This is not surprising since the optimal moving window is quite wide (700 samples). In the Ostracod case, the tracking model was again clearly the best, while the GWR model gave the poorest results (Fig. 6 and Table 3). In the two Stickleback fish cases, the WLS and GRW models gave very similar results (Figures 8 and 9, and Table 4), while the tracking model was much better.

It would, finally, be interesting to find more linear cases in the literature. It would be even more interesting to find cases with non-linear mean trait versus environment functions, as discussed in Ergon (2025). The tracking methodology used here can easily be extended to handle such cases. It would also be interesting to find more cases with obliquity cycles as vital parts of the environmental driver.

## Supporting information

MATAB Code

Data for Stickleback fish case

## Acknowledgements

I thank Mike Bell for information regarding the time span of the stickleback fish data and for the observation of possible obliquity cycles. I also thank University of South-Eastern Norway for support and funding.

## Data Availability Statement

Oxygen isotope data for Figures 4, 5 and 6 are archived as *Raw data for simulations* on bioRxiv, https://doi.org/10.1101/2024.10.30.621046

Oxygen isotope data for Figures 7, 8 and 9 are archived as *Data for Stickleback fish case* on bioRxiv, doi: https://doi.org/10.64898/2025.12.07.692820

## Appendix A Data for real data cases

**Table A1.** Data for AZ bryozoan case.

| Age<br>Ma | $\delta^{18}O(t)$<br>temperature | log AZ<br>area mean | SE<br>value |
| --- | --- | --- | --- |
| 2.1895 | 3.6518 | 11.3930 | 0.0500 |
| 1.9660 | 3.7409 | 11.4230 | 0.0500 |
| 1.8505 | 3.7487 | 11.4550 | 0.1200 |
| 0.9265 | 4.0233 | 11.6150 | 0.0500 |
| 0.6485 | 4.2053 | 11.6600 | 0.0400 |
| 0.5480 | 4.0763 | 11.5550 | 0.0450 |
| 0.5055 | 4.1545 | 11.6550 | 0.0400 |
| 0.4510 | 4.1775 | 11.7500 | 0.0850 |
| 0.3990 | 4.1735 | 11.7000 | 0.1200 |

**Table A2.** Data for OV bryozoan case.

| Age<br>Ma | $\delta^{18}O(t)$<br>temperature | log AZ<br>area mean | SE<br>value |
| --- | --- | --- | --- |
| 2.1895 | 3.6110 | 10.2600 | 0.0650 |
| 1.9660 | 3.6934 | 10.1700 | 0.0650 |
| 0.9265 | 4.0426 | 10.4300 | 0.0950 |
| 0.6485 | 4.0779 | 10.3900 | 0.0550 |
| 0.5480 | 4.0935 | 10.3600 | 0.0700 |
| 0.5055 | 4.1210 | 10.4300 | 0.0600 |
| 0.4510 | 4.1357 | 10.5000 | 0.1350 |

**Table A3.** Data for Ostracod case.

| Age<br>Ma | Mg/Ca<br>temperature | n | Mean valve length<br>$\mu m$ |
| --- | --- | --- | --- |
| 3.33 | 2.54 | 13 | 707 |
| 2.60 | 2.23 | 4 | 744 |
| 2.40 | 2.15 | 21 | 742 |
| 2.19 | 2.07 | 23 | 743 |
| 1.90 | 1.96 | 25 | 745 |
| 1.68 | 1.88 | 22 | 763 |
| 0.89 | 1.55 | 28 | 812 |
| 0.14 | 1.19 | 24 | 797 |
| 0.12 | 1.18 | 3 | 793 |
| 0.00 | 1.12 | 31 | 812 |

**Table A4.** Data for Stickleback fish case with 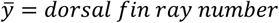.

| Time | N | $\bar{y}$ | SD |
| --- | --- | --- | --- |
| 0 | 30 | 8.8300 | 0.7470 |
| 0.0053 | 33 | 8.7600 | 0.6630 |
| 0.0100 | 30 | 8.8700 | 0.6810 |
| 0.0152 | 36 | 9.0600 | 0.7150 |
| 0.0193 | 55 | 8.5800 | 0.7620 |
| 0.0256 | 40 | 8.7500 | 0.8990 |
| 0.0305 | 75 | 8.5600 | 0.7750 |
| 0.0340 | 52 | 8.7700 | 0.7030 |
| 0.0366 | 22 | 9.1800 | 0.5010 |
| 0.0381 | 28 | 9.1800 | 0.7720 |
| 0.0410 | 54 | 8.9400 | 0.6270 |
| 0.0506 | 45 | 9.2000 | 0.5880 |
| 0.0552 | 35 | 9.1400 | 0.6480 |
| 0.0603 | 37 | 9.0500 | 0.6640 |
| 0.0653 | 41 | 9.1000 | 0.5390 |
| 0.0701 | 25 | 9.2000 | 0.7070 |
| 0.0751 | 30 | 9.1300 | 0.7300 |
| 0.0811 | 58 | 9.3100 | 0.7540 |
| 0.0865 | 32 | 9.7500 | 0.7180 |
| 0.0920 | 84 | 9.6300 | 0.8030 |
| 0.0934 | 82 | 10.0200 | 0.8460 |
| 0.0967 | 89 | 9.8000 | 0.8280 |
| 0.0994 | 95 | 9.6200 | 0.7320 |
| 0.1013 | 57 | 9.6700 | 0.7150 |
| 0.1070 | 45 | 9.8000 | 0.6940 |
| 0.1077 | 64 | 9.8400 | 0.7180 |

## Appendix B Obliquity cycles in Stickleback fish case

### B1. Data smoothing for feature extraction

For an elementary theoretical analysis, assume the model

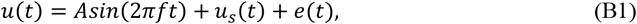

where *u*_*s*_(*t*) has a structure that is important for prediction of the mean traits in the block diagram in Fig. 1, while *e*(*t*) is unimportant high frequence noise. Also assume a sampling frequency *f*_*s*_ ≫ *f*, moving average smoothing with window sample size *n*, and *n* large enough to make the smoothed high frequency noise *e*_*smooth*_(*t*) ≈ 0.

For an integral number of periods in the moving average time window, the average of the sinusoidal component in Eq. (B1) is zero. For a non-integral number of periods we thus find 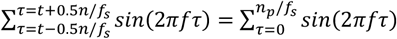, where 0 < *n*_*p*_ < *f*_*s*_/*f* (where *f*_*s*_/*f* is the number of samples per period). We thus find

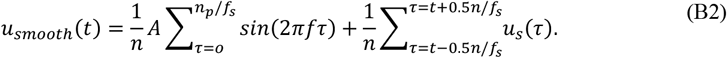

For a non-integral number of periods in the moving time window, the sinusoidal component will thus be seen in *u*_*smooth*_(*t*) with the frequency *f*, but with an amplitude depending on *n* and *n*_*p*_ according to Eq. (B2), while the structured information will be smoothed. The sampling window must thus be wide enough to filter out the noise component *e*(*t*), but small enough to retain the information on the sinusoidal component and the essential features in the structured information in *u*_*s*_(*t*). And the window should not be equal to an integer number of periods. As shown in Fig. 7, an optimal solution can be found by use of the window size as a tuning parameter.

As an example, Fig. B1, upper panel, shows the same smoothed mean trait data as in Fig. 7, upper panel, but detrended and over a longer time span. The mean trait apparently includes cycles with an approximate period of 41,000 years, i.e., with *f* = 24 *μHz*, which indicates variations caused by alterations in the Earth’s axial tilt (obliquity). Since *f*_*s*_ = 500 *μHz*, the number of samples per period is *f*_*s*_/*f* ≈ 20. From the theory above follows that the obliquity cycles should vanish with an integer number of cycles in the sampling window, for example for a window size of 164,000 years, as shown in Fig. B1, mid-panel. When the window size is further reduced, the obliquity cycles reappear, now with a larger amplitude owing to the reduced number of samples.

**Figure B1.**
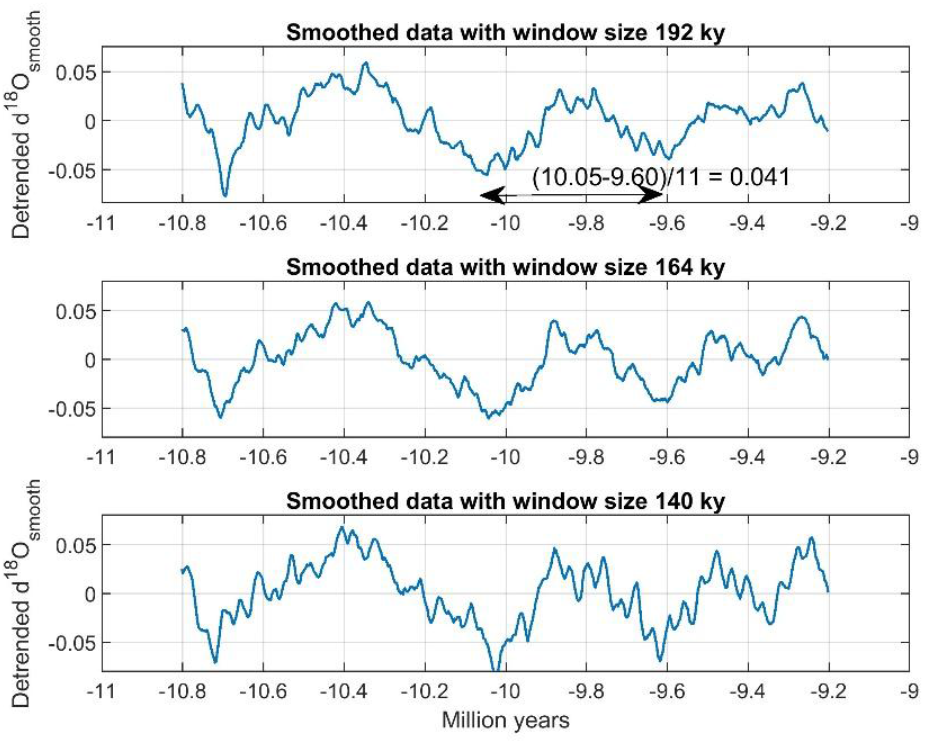
Obliquity cycles with an approximate period of 41,000 years, as seen with window size 192,000 years (upper panel). The cycles vanish with window size 164,000 years (mid-panel), and reappear with window size 140,000 years (lower panel).

Fig. B2 shows the smoothed mean trait data for different window sizes without detrending. With short window sizes the obliquity cycles are hidden in noise, with medium window sizes the cycles stand out, and with wide windows they are so much reduced that they again tend to be lost.

**Figure B2.**
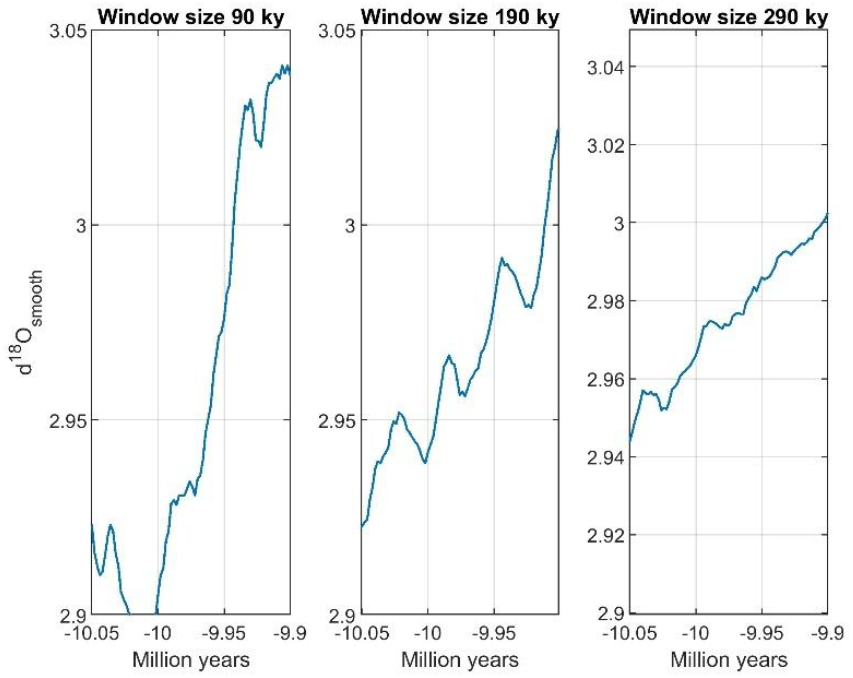
Smoothed *∂*^18^*O*(*t*) data for different time-window sizes.

### B2. Periodogram

In an attempt to find the obliquity cycles by spectral density analysis, the smoothed and detrended data *u*(*t*) in Fig. B1, upper panel, was used to find a periodogram, i.e., estimates of the power spectral density as function of frequency. First, a further smoothed signal *u*_*smooth*_(*t*) was found using a moving average time window of 80,000 years. Second, a residual *u*_*res*_(*t*) = *u*(*t*) − *u*_*smooth*_ was computed as shown in Fig. B3. This residual signal was finally used in the MATLAB function *periodogram*, with results as shown in Fig. B4, upper panel. Here, the peak at 27 cycles per million years (period 37,000 years) indicates the obliquity cycles. As shown in Fig. B1, upper panel, a more correct result is found by simply counting peaks over time. The peak at 51 cycles per million years (period 20,000 years) could possibly be explained by precession (axial wobble) with periods of 23,000 years (Milankovitch cycles - Wikipedia). The error in the estimated periods of the obliquity and precession cycles may thus be caused by interference between these cycles, as also indicated in Fig. B3 (red line), and possibly also other Milankovitch cycles. The periodogram with window size 164,000 years, such that the obliquity cycles in theory should vanish, is shown in Fig. B4, mid-panel. Here, the peak at 27 cycles per million years is very much reduced, but instead, and for unknown reasons, strong peaks at 21 and 32 cycles per million years are dominating features. It is interesting to note that the peak at 51 cycles per million years also is very much reduced with window size 164,000 years, and that 164,000 is close to the integer number 7 times the precession period of 23,000 years.

**Figure B3.**
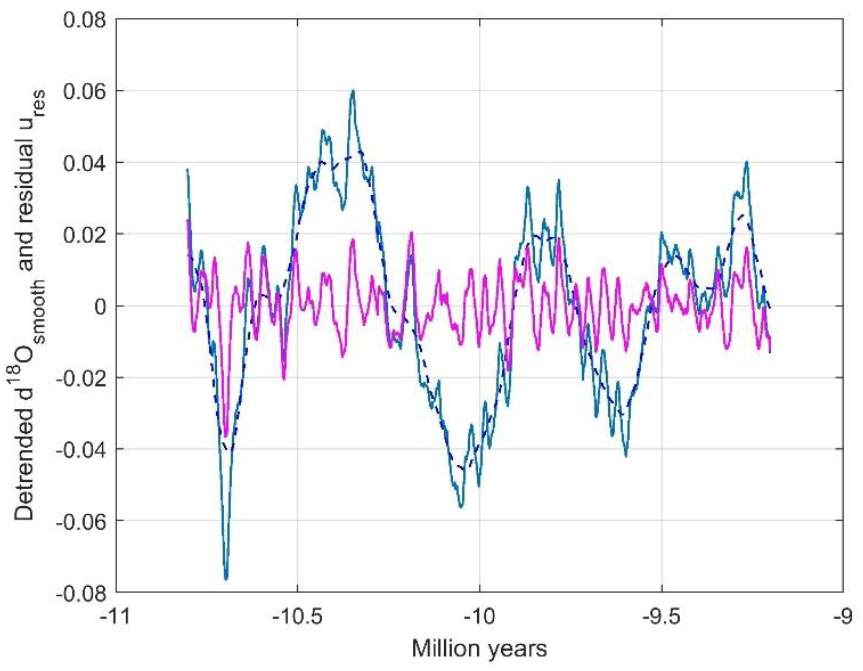
Detrended signal *u*(*t*) as in Fig, B1, upper panel, smoothed signal *u*_*smooth*_(*t*) (dashed blue line), and residual *u*_*res*_(*t*) = *u*(*t*) − *u*_*smooth*_(*t*) (red line).

**Figure B4.**
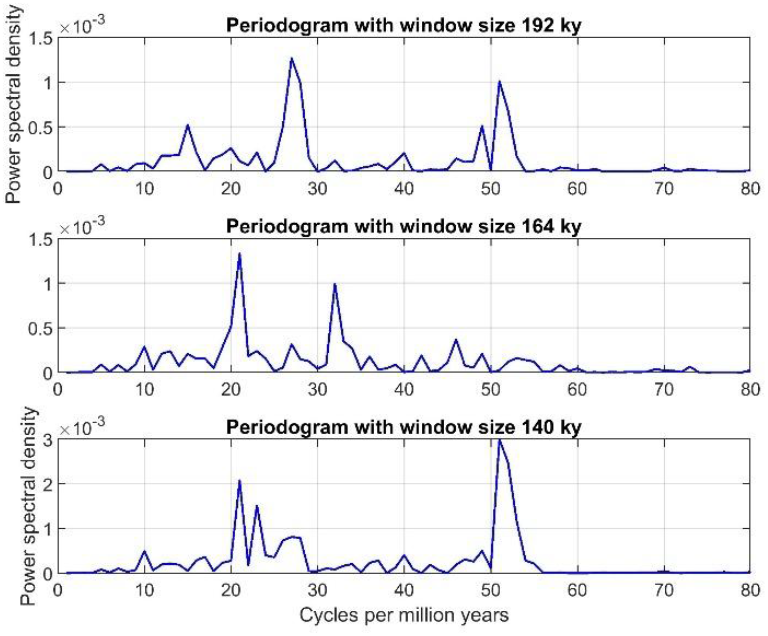
Periodograms corresponding to the time series in Fig. B1, as found from *u*_*res*_(*t*) as in Fig. B3 (red line) by use of the MATLAB function *periodogram*. Note the different scaling in the lower panel.

**Figure B4.**
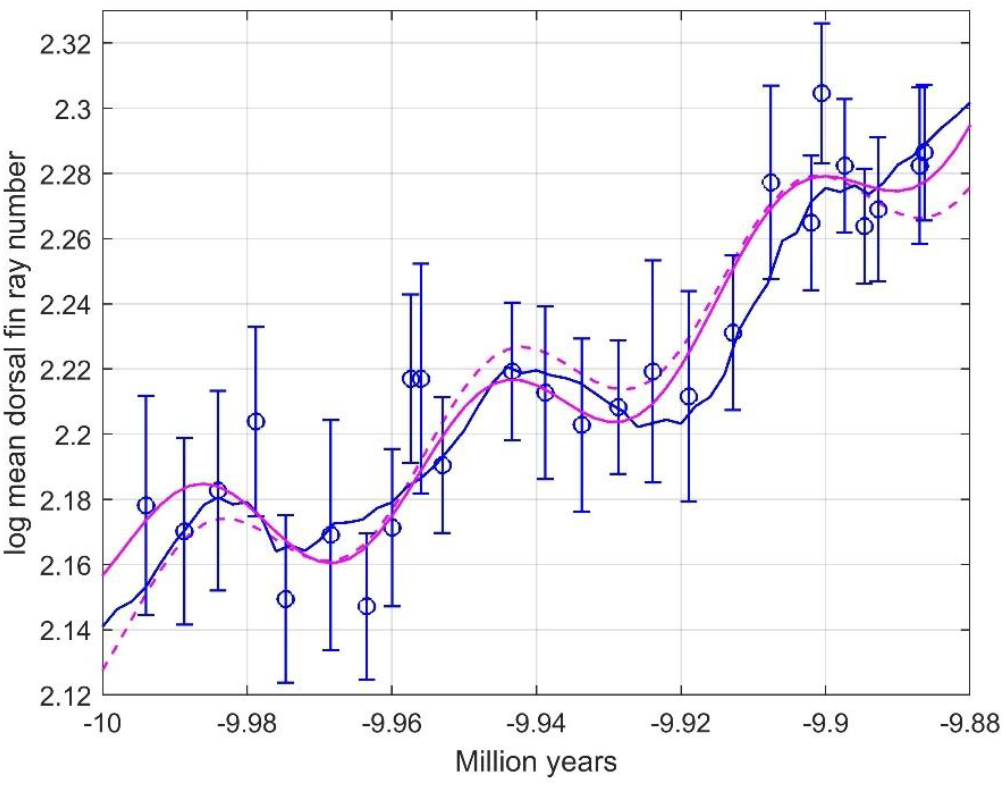
Tracking response as in Fig. 8 (blue line) and by use of Eq. (B3) with *b*_2_ as free variable (red line). The response with *b*_2_ = 0 is shown by dashed red line.

### B3. Sinusoidal model

For comparison with the tracking response in Fig. (8) we may use a non-linear prediction model with a sinus function,

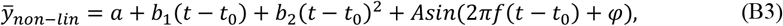

where 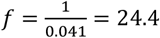 periods pr. million years. A numerical parameter search for minimum WMSE using the function *fmincon* in MATLAB resulted in *a* = 2.16, *b*_1_ = 0.27, *b*_2_ = 8.64, *A* = 0.0176 and *φ* = 0.52. With *b*_2_ = 0 this was changed into *a* = 2.14, *b*_1_ = 1.28, *b*_2_ = 0, *A* = 0.0177 and *φ* = 0.38. See Fig. B3 and Table B for results. Note that the model in Eq. (B3) with *b*_2_ as a free variable is only slightly better than the tracking model, while *b*_2_ = 0 gives a somewhat poorer model.

**Table B.** Prediction slope, WMSE and AIC results for Stickleback fish case.

| | Tracking model | Model in Eq. (B1) | Model in Eq. (B1) with $b_2 = 0$ |
| --- | --- | --- | --- |
| WMSE | 0.000216 | 0.000201 | 0.000258 |
| $AIC_c$ | -131 | -133 | -126 |
| Akaike weights | 0.28 | 0.72 | — |
| Akaike weights | 0.91 | — | 0.09 |

## Appendix C

MATLAB code for Stickleback fish case

clear

load Data_9800_to_10300.mat

Age=flipud(Age);

d18O=flipud(d18O);

tdata=-Age’; % Step size 0.002 My rawdata=d18O’;

window=96;

u=movmean(rawdata,window);

t0=-10+0.0060;

%% Sample data

tlabel1=[0 5317 9950 15182 19318 25551 30541 34030 36631 38052 41008 50629 55246];

tlabel2=[60289 65312 70119 75075 81150 86454 92006 93445 96659 99377 101306 106969 107667];

tlabel=[tlabel1 tlabel2]/1000000; tlabel=t0+tlabel;

n=[30 33 30 36 55 40 75 52 22 28 54 45 35 37 41 25 30 58 32 84 82 89 95 57 45 64];

y1=0.01*[883 876 887 906 858 875 856 877 918 918 894 920 914];

y2=0.01*[905 910 920 913 931 975 963 1002 980 962 967 980 984];

y=[y1 y2];

sd1=0.001*[747 663 681 715 762 899 775 703 501 772 627 588 648];

sd2=0.001*[664 539 707 730 754 718 803 846 828 732 715 694 718];

sd=[sd1 sd2]; var=sd.^2;

err0=sqrt(var./n);

ylabell=log(y);

for i=1:26

err(i)=err0(i)*(ylabell(i)/y(i))^1;

end

%% Tracking model

tplot=tdata;

ulabel=zeros(1,length(tlabel));

tlab=tlabel;

for i=1:length(tlabel)

for t=1:length(tplot)

if tplot(t)>min(tlab)

ulabel(1,i)=u(1,t);

tlab(1,i)=0;

break

end

end

end

w=err.^-2;

W=diag(w);

N=26;

k=3;

X=[ones(length(tlabel),1) ulabel’-2.9559*ones(length(tlabel),1)];

bls1=inv(X’*W*X)*X’*W*ylabell’

a1=bls1(1);

b1=bls1(2);

yhat1=a1+b1*(ulabel-2.9559*ones(1,26));

for t=1:length(u)

yhatplot(t)=a1+b1*(u(t)-2.9559);

end

WMSE_Tracking=sum((ylabell-yhat1)*W*(ylabell-yhat1)’)/sum(diag(W))

for j=1:N

lnw(j)=log(sqrt(1/w(j)));

z(j)=(ylabell(j)-yhat1(j)).^2*w(j);

end

AICc_Tracking=2*k+N*log(2*pi)+N+2*sum(lnw)+N*log(sum(z)/N)+2*k*(k+1)/(N-k-1);

%% WLS model

X=[ones(26,1) tlabel’];

bls=inv(X’*W*X)*X’*W*ylabell’;

a2=bls(1);

b2=bls(2);

for i=1:26

yhat2(i)=a2+b2*tlabel(i);

end

WMSE_WLS=(ylabell-yhat2)*W*(ylabell-yhat2)’/sum(diag(W))

for j=1:N

lnw(j)=log(sqrt(1/w(j)));

z(j)=(ylabell(j)-yhat2(j)).^2*w(j);

end

AICc_WLS=2*k+N*log(2*pi)+N+2*sum(lnw)+N*log(sum(z)/N)+2*k*(k+1)/(N-k-1);

%% GRW model

k=2;

N=25;

n_samples=ones(1,26);

var_samples=err.^2;

Tsample=tlabel;

for i=1:25

dT(i)=-Tsample(i)+Tsample(i+1);

dX(i)=ylabell(i+1)-ylabell(i);

nA(i)=n_samples(i);

nD(i)=n_samples(i+1);

varA(i)=var_samples(i);

varD(i)=var_samples(i+1);

end

% Contraints and

mustep_min=-10; mustep_max=10;

varstep_min=0; varstep_max=1;

par_lb=[mustep_min varstep_min];

par_ub=[mustep_max varstep_max];

par_guess=[0 0];

Aineq=[]; Bineq=[]; Aeq=[]; Beq=[];

fun_objective_handle=…

@(par)fun_objective(par,dT,dX,varA,nA,varD,nD);

[par_opt,fval,exitflag,output,lambda,grad,hessian] =…

fmincon(fun_objective_handle,par_guess,Aineq,Bineq,Aeq,Beq,par_lb,par_ub);

mustep=par_opt(1);

varstep=par_opt(2);

for i=1:25

varterm(i)=dT(i)*varstep+varA(i)/nA(i)+varD(i)/nD(i);

end

for i=1:25

Li(i)=-log(2*pi)/2-log(varterm(i))/2-(dX(i)-dT(i)*mustep)^2/(2*varterm(i));

end

L=sum(Li);

AICc_GRW=2*k-2*L+2*k*(k+1)/(N-k-1); X=ones(26,1);

b3=mustep;

a3=inv(X’*W*X)*X’*W*(ylabell’-b3*tlabel’); for i=1:26

yhat3(i)=a3+b3*tlabel(i);

end

WMSE_GRW=sum((ylabell-yhat3)*W*(ylabell-yhat3)’)/sum(diag(W));

% Akaike weights

D_Tracking=0;

D_WLS=AICc_WLS-AICc_Tracking;

D_GRW=AICc_GRW-AICc_Tracking;

L_WLS=exp(-0.5*D_WLS);

L_GRW=exp(-0.5*D_GRW);

Aweight_Tracking=1/(1+L_WLS+L_GRW);

Aweight_WLS=L_WLS/(1+L_WLS+L_GRW);

Aweight_GRW=L_GRW/(1+L_WLS+L_GRW);

yplot=a1+b1*(ulabel’-2.95*ones(26,1));

figure(7)

subplot(2,1,1)

plot(tplot,rawdata,’b’,’LineWidth’,1.0), hold on

plot(tplot,u,’m’,’LineWidth’,1.0), hold off, grid

axis([-10 -9.88 2.5 3.3])

xlabel(‘Million years’)

ylabel(‘d^1^8O and d^1^8O_s_m_o_o_t_h’)

subplot(2,1,2)

errorbar(ulabel,ylabell,err,’bo’,’LineWidth’,1.0), hold on, grid

plot(ulabel,yplot,’m’,’LineWidth’,1.0), hold off

axis([2.95 3.03 2.12 2.33])

xlabel(‘d^1^8O_s_m_o_o_t_h’)

ylabel(‘log mean dorsal fin ray number’)

figure(8)

errorbar(tlabel,ylabell,err,’bo’,’LineWidth’,1.0), hold on

plot(tplot,yhatplot,’b’,’LineWidth’,1.0)

plot(tlabel,yhat2,’m’);

plot(tlabel,yhat3,’m--’,’LineWidth’,1.0)

hold off, grid

axis([-10 -9.88 2.12 2.33])

xlabel(‘Million years’)

ylabel(‘log mean dorsal fin ray number’)

b2

b3

WMSE_Tracking

AICc_Tracking

WMSE_WLS

AICc_WLS

WMSE_GRW AICc_GRW

Aweight_Tracking

Aweight_WLS

Aweight_GRW

%% Parameter search

function f = fun_objective(par,dT,dX,varA,nA,varD,nD)

mustep=par(1);

varstep=par(2);

for i=1:25

varterm=dT(i)*varstep+varA(i)/nA(i)+varD(i)/nD(i);

Li(i)=-log(2*pi)/2-log(varterm)/2-(dX(i)-dT(i)*mustep)^2/(2*varterm); end

L=sum(Li);

f=-L;

end

