## Supplementary material for "Examples of adaptive peak tracking as found in the fossil record": MATAB Code

### MATLAB code for

Rolf Ergon

University of South-Eastern Norway

December 7, 2025

##### Figures 4 and 5

```
clear

load Data_9800_to_10300.mat

Age=flipud(Age);
d18O=flipud(d18O);

tdata=-Age';        % Step size 0.002 My
udata=d18O';

window=96;
ufilt=movmean(udata,window);
t0=-10+0.006;

%% Sample data
tlabel1=[0 5317 9950 15182 19318 25551 30541 34030 36631 38052 41008 50629 55246];
tlabel2=[60289 65312 70119 75075 81150 86454 92006 93445 96659 99377 101306 106969 107667];
tlabel=[tlabel1 tlabel2]/1000000;
tlabel=t0+tlabel;
n=[30 33 30 36 55 40 75 52 22 28 54 45 35 37 41 25 30 58 32 84 82 89 95 57 45 64];
y1=0.01*[883 876 887 906 858 875 856 877 918 918 894 920 914 ];
y2=0.01*[905 910 920 913 931 975 963 1002 980 962 967 980 984];
y=[y1 y2];
sd1=0.001*[747 663 681 715 762 899 775 703 501 772 627 588 648  ];
sd2=0.001*[664 539 707 730 754 718 803 846 828 732 715 694 718];
sd=[sd1 sd2];
var=sd.^2;
err0=sqrt(var./n);
```

```
ylabell=log(y);

for i=1:26

   err(i)=err0(i)*(ylabell(i)/y(i))^1;

end


%% Tracking model

tplot=tdata;

ulabel=zeros(1,length(tlabel));

ufilt=ufilt;

tlab=tlabel;

for i=1:length(tlabel)

   for t=1:length(tplot)

      if tplot(t)>min(tlab)

         ulabel(1,i)=ufilt(1,t);

         tlab(1,i)=0;

         break

      end

   end

end


w=err.^-2;

W=diag(w);


N=26;

k=3;


X=[ones(length(tlabel),1) ulabel'-2.9559*ones(length(tlabel),1)];

bls1=inv(X'*W*X)*X'*W*ylabell'

a1=bls1(1);

b1=bls1(2);


yhat1=a1+b1*(ulabel-2.9559*ones(1,26));


for t=1:length(ufilt)

   yhatplot(t)=a1+b1*(ufilt(t)-2.9559);

end


WMSE_Tracking=sum((ylabell-yhat1)*W*(ylabell-yhat1)')/sum(diag(W))
```

```
for j=1:N
  lnw(j)=log(sqrt(1/w(j)));
  z(j)=(ylabell(j)-yhat1(j)).^2*w(j);
end

AICc_Tracking=2*k+N*log(2*pi)+N+2*sum(lnw)+N*log(sum(z)/N)+2*k*(k+1)/(N-k-1);

%% WLS model
X=[ones(26,1) tlabel'];
bls=inv(X'*W*X)*X'*W*ylabell';
a2=bls(1);
b2=bls(2);
for i=1:26
   yhat2(i)=a2+b2*tlabel(i);
end

WMSE_WLS=(ylabell-yhat2)*W*(ylabell-yhat2)'/sum(diag(W))

for j=1:N
  lnw(j)=log(sqrt(1/w(j)));
  z(j)=(ylabell(j)-yhat2(j)).^2*w(j);
end

AICc_WLS=2*k+N*log(2*pi)+N+2*sum(lnw)+N*log(sum(z)/N)+2*k*(k+1)/(N-k-1);

%% GRW model
k=2;
N=25;
n_samples=ones(1,26);
var_samples=err.^2;

c=1;
Tsample=c*tlabel;

for i=1:25
   dT(i)=-Tsample(i)+Tsample(i+1);
   dX(i)=ylabell(i+1)-ylabell(i);
   nA(i)=n_samples(i);
```

```
   nD(i)=n_samples(i+1);
   varA(i)=var_samples(i);
   varD(i)=var_samples(i+1);
end

% Contraints
mustep_min=-10; mustep_max=10;
varstep_min=0; varstep_max=1;

par_lb=[mustep_min varstep_min];
par_ub=[mustep_max varstep_max];

par_guess=[0 0];
Aineq=[]; Bineq=[]; Aeq=[]; Beq=[];
fun_objective_handle=...
   @(par)fun_objective(par,dT,dX,varA,nA,varD,nD);
[par_opt,fval,exitflag,output,lambda,grad,hessian] =...
fmincon(fun_objective_handle,par_guess,Aineq,Bineq,Aeq,Beq,par_lb,par_ub);
output;

mustep=par_opt(1);
varstep=par_opt(2);

for i=1:25
varterm(i)=dT(i)*varstep+varA(i)/nA(i)+varD(i)/nD(i);
end

for i=1:25
   Li(i)=-log(2*pi)/2-log(varterm(i))/2-(dX(i)-dT(i)*mustep)^2/(2*varterm(i));
end

L=sum(Li);
AICc_GRW=2*k-2*L+2*k*(k+1)/(N-k-1);

X=ones(26,1);
b3=c*mustep;
a3=inv(X'*W*X)*X'*W*(ylabell'-b3*tlabel');
for i=1:26
   yhat3(i)=a3+b3*tlabel(i);
```

```
end

WMSE_GRW=sum((ylabell-yhat3)*W*(ylabell-yhat3)')/sum(diag(W));

yplot=a1+b1*(ulabel'-2.95*ones(26,1));

figure(4)
subplot(2,1,1)
plot(tplot,udata,'b','LineWidth',1.0), hold on
plot(tplot,ufilt,'m','LineWidth',1.0), hold off, grid
axis([-10 -9.88 2.5 3.3])
xlabel('Million years')
ylabel('d^1^8O and d^1^8O_f_i_l_t')

subplot(2,1,2)
errorbar(ulabel,ylabell,err,'bo','LineWidth',1.0), hold on, grid
plot(ulabel,yplot,'m','LineWidth',1.0), hold off
% axis([-10 -9.88 2.9 3.07])
xlabel('d^1^8O_f_i_l_t')
ylabel('log mean trait')

figure(5)
errorbar(tlabel,ylabell,err,'bo','LineWidth',1.0), hold on
plot(tplot,yhatplot,'b','LineWidth',1.0)
plot(tlabel,yhat2,'m');
plot(tlabel,yhat3,'m--','LineWidth',1.0)
hold off, grid
axis([-10 -9.88 2.12 2.33])
xlabel('Million years')
ylabel('log mean body size')

b2
b3
WMSE_Tracking
AICc_Tracking
WMSE_WLS
AICc_WLS
WMSE_GRW
AICc_GRW
```

```
%% Parameter search
function f = fun_objective(par,dT,dX,varA,nA,varD,nD)
mustep=par(1);
varstep=par(2);

for i=1:25
   varterm=dT(i)*varstep+varA(i)/nA(i)+varD(i)/nD(i);
   Li(i)=-log(2*pi)/2-log(varterm)/2-(dX(i)-dT(i)*mustep)^2/(2*varterm);
end

L=sum(Li);

f=-L

end
```

#### Figure 6

```
clear
s2=467;              % micrometer^2

window=80;

%% Load O18 data, order and plot
load O18.mat
x=5320-x;
x=flipud(x);
u=flipud(y);

for t=2:length(x)
   dx(t)=x(t)-x(t-1);
end

x=0.001*x;             % Million year scale
ufilt=movmean(u,window);
ufilt1=movmean(u,0.4*window);
ufilt2=movmean(u,0.5*window);
ufilt3=movmean(u,1*window);
ufilt=[ufilt1(1:329+736) ; ufilt2(330+736:779+736) ; ufilt3(780+736:2115)];
```

```
tplot=x-5.3-0.02;          % -0.02 for zero point correction

%% Sample data
Data=[
3.33 1,962 2.54 13 707
2.60 3,427 2.23 4 744
2.40 1,253 2.15 21 742
2.19 1,253 2.07 23 743
1.90 1,253 1.96 25 745
1.68 1,253 1.88 22 763
0.89 2,873 1.55 28 812
0.14 2,873 1.19 24 797
0.12 3,427 1.18 3 793
0.00 2,781 1.12 31 812
];

tlabel=-Data(:,1)';
y=Data(:,6)';
ylabell=log(Data(:,6))';
n=Data(:,5)';

for i=1:10
   err(i)=sqrt(s2/n(i))*ylabell(i)/y(i);
end

%% Tracking model
ulabel=zeros(1,length(tlabel));
ufilt=ufilt';
tlab=tlabel;
for i=1:length(tlabel)
   for t=1:length(tplot)
      if tplot(t)>min(tlab)
         ulabel(1,i)=ufilt(1,t)
         tlab(1,i)=0;
         break
      end
   end
end
```

```
var=err.^2;
W1=diag(var);
W1=inv(W1);
w=1./var;

X=[ones(length(tlabel),1) ulabel'-3.15*ones(length(tlabel),1)];
bls1=inv(X'*W1*X)*X'*W1*ylabell'
a1=bls1(1);
b1=bls1(2);
for i=1:10
  yhat1(i)=a1+b1*(ulabel(i)-3.15);
end

for t=1:2115
  yhatplot(t)=a1+b1*(ufilt(t)-3.15);
end

WMSE_Tracking=sum((ylabell-yhat1)*W1*(ylabell-yhat1)')/sum(diag(W1));

N=10;
k=3;

for i=1:N
  lnw(i)=log(sqrt(1/w(i)));
  z(i)=(ylabell(i)-yhat1(i)).^2*w(i);
end

AICc_Tracking=2*k+N*log(2*pi)+N+2*sum(lnw)+N*log(sum(z)/N)+2*k*(k+1)/(N-k-1);

%% WLS model
W2=W1;

X=[ones(10,1) tlabel'];
bls=inv(X'*W2*X)*X'*W2*ylabell';
a2=bls(1);
b2=bls(2);
for i=1:10
  yhat2(i)=a2+b2*tlabel(i);
```

```
end

WMSE_WLS=(ylabell-yhat2)*W2*(ylabell-yhat2)'/sum(diag(W2));

for j=1:N
  lnw(j)=log(sqrt(1/w(j)));
  z(j)=(ylabell(j)-yhat2(j)).^2*w(j);
end

AICc_WLS=2*k+N*log(2*pi)+N+2*sum(lnw)+N*log(sum(z)/N)+2*k*(k+1)/(N-k-1);

%% GRW model
k=2;
N=9;
W3=W2;
n_samples=ones(1,10);
var_samples=err.^2;

c=1;
Tsample=c*tlabel;

for i=1:9
   dT(i)=-Tsample(i)+Tsample(i+1);
   dX(i)=ylabell(i+1)-ylabell(i);
   nA(i)=n_samples(i);
   nD(i)=n_samples(i+1);
   varA(i)=var_samples(i);
   varD(i)=var_samples(i+1);
end

% Contraints
mustep_min=-1; mustep_max=1;
varstep_min=0; varstep_max=1;

par_lb=[mustep_min varstep_min];
par_ub=[mustep_max varstep_max];

par_guess=[0 0];
Aineq=[]; Bineq=[]; Aeq=[]; Beq=[];
```

```
fun_objective_handle=...
   @(par)fun_objective(par,dT,dX,varA,nA,varD,nD);
[par_opt,fval,exitflag,output,lambda,grad,hessian] =...
fmincon(fun_objective_handle,par_guess,Aineq,Bineq,Aeq,Beq,par_lb,par_ub);
output;

mustep=par_opt(1);
varstep=par_opt(2);

for i=1:9
varterm(i)=dT(i)*varstep+varA(i)/nA(i)+varD(i)/nD(i);
end

for i=1:9
   Li(i)=-log(2*pi)/2-log(varterm(i))/2-(dX(i)-dT(i)*mustep)^2/(2*varterm(i));
end

L=sum(Li);
AICc_GRW=2*k-2*L+2*k*(k+1)/(N-k-1);

X=ones(10,1);
b3=c*mustep;
a3=inv(X'*W3*X)*X'*W3*(ylabell'-b3*tlabel');
for i=1:10
   yhat3(i)=a3+b3*tlabel(i);
end

WMSE_GRW=sum((ylabell-yhat3)*W3*(ylabell-yhat3)')/sum(diag(W3));

figure(6)
errorbar(tlabel,ylabell,err,'bo','LineWidth',1.0), hold on
plot(tplot,yhatplot,'b','LineWidth',1.0)
plot(tlabel,yhat2,'m','LineWidth',1.0)
plot(tlabel,yhat3,'m--','LineWidth',1.0)
hold off, grid
axis([-3.4 0.1 6.5 6.8])
xlabel('Million years')
ylabel('log mean body size')
```

```
b2
b3
WMSE_Tracking
AICc_Tracking
WMSE_WLS
AICc_WLS
WMSE_GRW
AICc_GRW

%% Parameter search
function f = fun_objective(par,dT,dX,varA,nA,varD,nD)
mustep=par(1);
varstep=par(2);

for i=1:9
   varterm=dT(i)*varstep+varA(i)/nA(i)+varD(i)/nD(i);
   Li(i)=-log(2*pi)/2-log(varterm)/2-(dX(i)-dT(i)*mustep)^2/(2*varterm);
end

L=sum(Li);

f=-L

end
```

#### Figure 7

```
clear
window=700;

%% Load O18 data, order and plot
load O18.mat
x=5320-x;
x=flipud(x);
u=flipud(y);

for t=2:length(x)
   dx(t)=x(t)-x(t-1);
end
```

```
x=0.001*x;              % Million year scale
ufilt=movmean(u,window);
ufilt=ufilt(737:2115);
x=x(737:2115);
u=u(737:2115);

tplot=x-5.3-0.02;        % -0.02 for zero point correction

% Sample data from Liow et al. (2024)
tlabel=[-2.1895 -1.9660 -0.9265 -0.6485 -0.5480 -0.5055 -0.4510 ];
ylabell=[10.26 10.17 10.43 10.39 10.36 10.43 10.50 ];
err=[0.065 0.065 0.095 0.055 0.07 0.06 0.135 ];

%% Tracking model
ulabel=[mean(ufilt(54))
    mean(ufilt(143))
    mean(ufilt(616))
    mean(ufilt(755))
    mean(ufilt(831))
    mean(ufilt(874))
    mean(ufilt(928))
    ]';
w=err.^-2;
W=diag(w);
W1=W;
X=[ones(7,1) ulabel'-3.6*ones(7,1)];
bls=inv(X'*W1*X)*X'*W1*ylabell';
ahat1=bls(1);
bhat1=bls(2);

for t=1:1379
   yhatplot(t)=ahat1+bhat1*(ufilt(t)-3.6);
end
yhat1=ahat1+bhat1*(ulabel-3.6*ones(1,7));

WMSE_Tracking=sum((ylabell-yhat1)*W1*(ylabell-yhat1)')/sum(diag(W1));

N=7;
k=3
```

```
for i=1:N
   lnw(i)=log(sqrt(1/w(i)));
   z(i)=(ylabell(i)-yhat1(i)).^2*w(i);
end

AICc_Tracking=2*k+N*log(2*pi)+N+2*sum(lnw)+N*log(sum(z)/N)+2*k*(k+1)/(N-k-1);



%% WLS model
W2=W1;

X=[ones(7,1) tlabel'];
bls=inv(X'*W2*X)*X'*W2*ylabell';
ahat2=bls(1);
bhat2=bls(2);
for i=1:7
    yhat2(1,i)=ahat2+bhat2*tlabel(i);
end

WMSE_WLS=sum((ylabell-yhat2)*W2*(ylabell-yhat2)')/sum(diag(W2));

for i=1:N
   lnw(i)=log(sqrt(1/w(i)));
   z(i)=(ylabell(i)-yhat2(i)).^2*w(i);
end

AICc_WLS=2*k+N*log(2*pi)+N+2*sum(lnw)+N*log(sum(z)/N)+2*k*(k+1)/(N-k-1);



%% GRW model
W3=W2;
k=2;
N=6;

n_samples=ones(1,7);
var_samples=err.^2;

c=1;
```

```
Tsample=c*tlabel;

for i=1:6
   dT(i)=-Tsample(i)+Tsample(i+1);
   dX(i)=ylabell(i+1)-ylabell(i);
   nA(i)=n_samples(i);
   nD(i)=n_samples(i+1);
   varA(i)=var_samples(i);
   varD(i)=var_samples(i+1);
end

% Contraints
mustep_min=-1; mustep_max=1;
varstep_min=0; varstep_max=1;

par_lb=[mustep_min varstep_min];
par_ub=[mustep_max varstep_max];

par_guess=[0 0];
Aineq=[]; Bineq=[]; Aeq=[]; Beq=[];
fun_objective_handle=...
   @(par)fun_objective(par,dT,dX,varA,nA,varD,nD);
[par_opt,fval,exitflag,output,lambda,grad,hessian] =...
fmincon(fun_objective_handle,par_guess,Aineq,Bineq,Aeq,Beq,par_lb,par_ub);

par_opt;
varstep=par_opt(2)
mustep=par_opt(1)

for i=1:6
varterm(i)=dT(i)*varstep+varA(i)/nA(i)+varD(i)/nD(i);
   Li(i)=-log(2*pi)/2-log(varterm(i))/2-(dX(i)-dT(i)*mustep)^2/(2*varterm(i));
end

L=sum(Li);
AICc_GRW=2*k-2*L+2*k*(k+1)/(N-k-1)

X=ones(7,1);
bhat3=c*mustep;
```

```
ahat3=inv(X'*W3*X)*X'*W3*(ylabell'-bhat3*tlabel');
for i=1:7
   yhat3(i)=ahat3+bhat3*tlabel(i);
end

WMSE_GRW=sum((ylabell-yhat3)*W3*(ylabell-yhat3)')/sum(diag(W3));

bhat2
bhat3
WMSE_Tracking
AICc_Tracking
WMSE_WLS
AICc_WLS
WMSE_GRW
AICc_GRW

figure(7)
errorbar(tlabel,ylabell,err,'bo','LineWidth',1.0), hold on
plot(tplot,yhatplot,'b','LineWidth',1.0)
plot(tlabel,yhat2,'m','LineWidth',1.0)
plot(tlabel,yhat3,'--m','LineWidth',1.0)
hold off, grid
axis([-2.35 -0.3 10.1 10.6])
xlabel('Million years')
ylabel('log OV area mean')

%% Parameter search
function f = fun_objective(par,dT,dX,varA,nA,varD,nD)
mustep=par(1);
varstep=par(2);

for i=1:6
   varterm(i)=dT(i)*varstep+varA(i)/nA(i)+varD(i)/nD(i);
   Li(i)=-log(2*pi)/2-log(varterm(i))/2-(dX(i)-dT(i)*mustep)^2/(2*varterm(i));
end

L=-sum(Li);

f=L;
```

```
end
```

#### Figure 8

```
clear
window=100;

%% Load O18 data, order and plot
load O18.mat
x=5320-x;
x=flipud(x);
u=flipud(y);

x=0.001*x;              % Million year scale
ufilt=movmean(u,window);
x=x(737:2115);
u=u(737:2115);
ufilt=ufilt(737:2115);
tplot=x-5.3-0.02;       % -0.2 for zero point correction

% Sample data from Liow et al. (2024)
tlabel=[-2.1895 -1.9660 -1.8505 -0.9265 -0.6485 -0.5480 -0.5055 -0.4510 -0.3990 ];
ylabell=[11.393 11.423 11.455 11.615 11.66 11.555 11.655 11.75 11.7 ];
err=[0.05 0.05 0.12 0.05 0.04 0.045 0.04 0.085 0.12];

%% Tracking model
ulabel=[mean(ufilt(54))
    mean(ufilt(143))
    mean(ufilt(189))
    mean(ufilt(616))
    mean(ufilt(755))
    mean(ufilt(831))
    mean(ufilt(874))
    mean(ufilt(928))
    mean(ufilt(980))
    ]';
w=err.^-2;
W=diag(w);
W1=W;
```

```
X=[ones(9,1) ulabel'-3.6*ones(9,1)];
bls=inv(X'*W1*X)*X'*W1*ylabell';
ahat1=bls(1);
bhat1=bls(2);

for t=1:1379
   yhatplot(t)=ahat1+bhat1*(ufilt(t)-3.6);
end
yhat1=ahat1+bhat1*(ulabel-3.6*ones(1,9));

WMSE_Tracking=sum((ylabell-yhat1)*W1*(ylabell-yhat1)')/sum(diag(W1));

N=9;
k=3

for i=1:N
  lnw(i)=log(sqrt(1/w(i)));
  z(i)=(ylabell(i)-yhat1(i)).^2*w(i);
end

AICc_Tracking=2*k+N*log(2*pi)+N+2*sum(lnw)+N*log(sum(z)/N)+2*k*(k+1)/(N-k-1);

%% WLS model
W2=W1;

X=[ones(9,1) tlabel'];
bls=inv(X'*W2*X)*X'*W2*ylabell';
ahat2=bls(1);
bhat2=bls(2);
for i=1:9
   yhat2(1,i)=ahat2+bhat2*tlabel(i);
end

WMSE_WLS=sum((ylabell-yhat2)*W2*(ylabell-yhat2)')/sum(diag(W2));

for i=1:N
  lnw(i)=log(sqrt(1/w(i)));
  z(i)=(ylabell(i)-yhat2(i)).^2*w(i);
end
```

```
AICc_WLS=2*k+N*log(2*pi)+N+2*sum(lnw)+N*log(sum(z)/N)+2*k*(k+1)/(N-k-1);

%% GRW model
W3=W2;
k=2;
N=8;

n_samples=ones(1,10);
var_samples=err.^2;

c=1;
Tsample=c*tlabel;

for i=1:8
   dT(i)=-Tsample(i)+Tsample(i+1);
   dX(i)=ylabell(i+1)-ylabell(i);
   nA(i)=n_samples(i);
   nD(i)=n_samples(i+1);
   varA(i)=var_samples(i);
   varD(i)=var_samples(i+1);
end

% Contraints
mustep_min=-1; mustep_max=1;
varstep_min=0; varstep_max=1;

par_lb=[mustep_min varstep_min];
par_ub=[mustep_max varstep_max];

par_guess=[0 0];
Aineq=[]; Bineq=[]; Aeq=[]; Beq=[];
fun_objective_handle=...
   @(par)fun_objective(par,dT,dX,varA,nA,varD,nD);
[par_opt,fval,exitflag,output,lambda,grad,hessian] =...
fmincon(fun_objective_handle,par_guess,Aineq,Bineq,Aeq,Beq,par_lb,par_ub);

par_opt;
varstep=par_opt(2)
```

```
mustep=par_opt(1)

for i=1:8
varterm(i)=dT(i)*varstep+varA(i)/nA(i)+varD(i)/nD(i);
   Li(i)=-log(2*pi)/2-log(varterm(i))/2-(dX(i)-dT(i)*mustep)^2/(2*varterm(i));
end

L=sum(Li);
AICc_GRW=2*k-2*L+2*k*(k+1)/(N-k-1)

X=ones(9,1);
bhat3=c*mustep;
ahat3=inv(X'*W3*X)*X'*W3*(ylabell'-bhat3*tlabel');
for i=1:9
   yhat3(i)=ahat3+bhat3*tlabel(i);
end

WMSE_GRW=sum((ylabell-yhat3)*W3*(ylabell-yhat3)')/sum(diag(W3));

bhat2
bhat3
WMSE_Tracking
AICc_Tracking
WMSE_WLS
AICc_WLS
WMSE_GRW
AICc_GRW

figure(8)
errorbar(tlabel,ylabell,err,'bo','LineWidth',1.0), hold on
plot(tplot,yhatplot,'b','LineWidth',1.0)
plot(tlabel,yhat2,'m','LineWidth',1.0)
plot(tlabel,yhat3,'--m','LineWidth',1.0)
hold off, grid
axis([-2.35 -0.3 11.3 11.85])
xlabel('Million years')
ylabel('log AZ area mean')

%% Parameter search
```

```
function f = fun_objective(par,dT,dX,varA,nA,varD,nD)

mustep=par(1);

varstep=par(2);


for i=1:8

   varterm(i)=dT(i)*varstep+varA(i)/nA(i)+varD(i)/nD(i);

   Li(i)=-log(2*pi)/2-log(varterm(i))/2-(dX(i)-dT(i)*mustep)^2/(2*varterm(i));

end


L=-sum(Li);


f=L;


end
```
