## Supplementary material for "Examples of adaptive peak tracking as found in the fossil record": Data for Stickleback fish case

### Data for use in the Stickleback fish case

Data_9800_to_10300 =

| | |
|---|---|
| 9.8000 | 2.8800 |
| 9.8020 | 3.0100 |
| 9.8040 | 2.9200 |
| 9.8060 | 3.0950 |
| 9.8080 | 3.2700 |
| 9.8100 | 2.9600 |
| 9.8120 | 3.4600 |
| 9.8140 | 3.2600 |
| 9.8160 | 3.3100 |
| 9.8180 | 3.5150 |
| 9.8200 | 3.1000 |
| 9.8220 | 3.0960 |
| 9.8240 | 3.0920 |
| 9.8260 | 3.0880 |
| 9.8280 | 3.0840 |
| 9.8300 | 3.0800 |
| 9.8320 | 2.8200 |
| 9.8340 | 2.9500 |
| 9.8360 | 2.9000 |
| 9.8380 | 2.8900 |
| 9.8400 | 2.9650 |
| 9.8420 | 2.9900 |
| 9.8440 | 2.9100 |
| 9.8460 | 3.0400 |
| 9.8480 | 2.8000 |
| 9.8500 | 3.1600 |
| 9.8520 | 3.2650 |
| 9.8540 | 3.2700 |
| 9.8560 | 3.2100 |
| 9.8580 | 3.1900 |
| 9.8600 | 3.1400 |
| 9.8620 | 3.1000 |
| 9.8640 | 3.0250 |
| 9.8660 | 3.1100 |
| 9.8680 | 3.0700 |
| 9.8700 | 2.9900 |
| 9.8720 | 3.0300 |
| 9.8740 | 3.2200 |
| 9.8760 | 3.1000 |
| 9.8780 | 2.7900 |
| 9.8800 | 2.9800 |
| 9.8820 | 2.4800 |
| 9.8840 | 2.7300 |
| 9.8860 | 3.0000 |
| 9.8880 | 2.7600 |
| 9.8900 | 3.0500 |
| 9.8920 | 3.0300 |

| | |
|---|---|
| 9.8940 | 3.1200 |
| 9.8960 | 3.2000 |
| 9.8980 | 3.2400 |
| 9.9000 | 3.2300 |
| 9.9020 | 3.1400 |
| 9.9040 | 3.1800 |
| 9.9060 | 3.1400 |
| 9.9080 | 2.9000 |
| 9.9100 | 2.9200 |
| 9.9120 | 3.0200 |
| 9.9140 | 3.0400 |
| 9.9160 | 3.0200 |
| 9.9180 | 3.0100 |
| 9.9200 | 3.0100 |
| 9.9220 | 2.8700 |
| 9.9240 | 2.9700 |
| 9.9260 | 2.9600 |
| 9.9280 | 3.0600 |
| 9.9300 | 2.9600 |
| 9.9320 | 3.0900 |
| 9.9340 | 3.1900 |
| 9.9360 | 3.1000 |
| 9.9380 | 3.0900 |
| 9.9400 | 3.1800 |
| 9.9420 | 3.0000 |
| 9.9440 | 3.0800 |
| 9.9460 | 3.3200 |
| 9.9480 | 3.1000 |
| 9.9500 | 3.2300 |
| 9.9520 | 2.9500 |
| 9.9540 | 3.0800 |
| 9.9560 | 3.0700 |
| 9.9580 | 3.0100 |
| 9.9600 | 2.9900 |
| 9.9620 | 2.8900 |
| 9.9640 | 2.8800 |
| 9.9660 | 2.8400 |
| 9.9680 | 2.8600 |
| 9.9700 | 2.9850 |
| 9.9720 | 2.7900 |
| 9.9740 | 2.8900 |
| 9.9760 | 2.8800 |
| 9.9780 | 2.8100 |
| 9.9800 | 2.8300 |
| 9.9820 | 2.7800 |
| 9.9840 | 2.7900 |
| 9.9860 | 2.8800 |
| 9.9880 | 2.7000 |
| 9.9900 | 2.8200 |
| 9.9920 | 3.0400 |
| 9.9940 | 2.8900 |

| | |
|---|---|
| 9.9960 | 2.9900 |
| 9.9980 | 2.8450 |
| 10.0000 | 2.7000 |
| 10.0020 | 2.8100 |
| 10.0040 | 2.6800 |
| 10.0060 | 2.8600 |
| 10.0080 | 2.8500 |
| 10.0100 | 2.7000 |
| 10.0120 | 2.6900 |
| 10.0140 | 2.9300 |
| 10.0160 | 2.7700 |
| 10.0180 | 3.1600 |
| 10.0200 | 3.0200 |
| 10.0220 | 3.0100 |
| 10.0240 | 3.1100 |
| 10.0260 | 3.1000 |
| 10.0280 | 3.0900 |
| 10.0300 | 3.0700 |
| 10.0320 | 3.0550 |
| 10.0340 | 3.0400 |
| 10.0360 | 3.0000 |
| 10.0380 | 2.9900 |
| 10.0400 | 2.9100 |
| 10.0420 | 2.8770 |
| 10.0440 | 2.8430 |
| 10.0460 | 2.8100 |
| 10.0480 | 2.8800 |
| 10.0500 | 2.8800 |
| 10.0520 | 2.9050 |
| 10.0540 | 2.9300 |
| 10.0560 | 2.7500 |
| 10.0580 | 3.0100 |
| 10.0600 | 2.8600 |
| 10.0620 | 2.9350 |
| 10.0640 | 3.0100 |
| 10.0660 | 2.9100 |
| 10.0680 | 2.9100 |
| 10.0700 | 2.8900 |
| 10.0720 | 2.8700 |
| 10.0740 | 3.0900 |
| 10.0760 | 3.0230 |
| 10.0780 | 2.9570 |
| 10.0800 | 2.8900 |
| 10.0820 | 2.9100 |
| 10.0840 | 2.6700 |
| 10.0860 | 2.8000 |
| 10.0880 | 2.8050 |
| 10.0900 | 2.8100 |
| 10.0920 | 2.9600 |
| 10.0940 | 3.0000 |
| 10.0960 | 2.8600 |

| | |
|---|---|
| 10.0980 | 2.9900 |
| 10.1000 | 3.1200 |
| 10.1020 | 3.1800 |
| 10.1040 | 3.1200 |
| 10.1060 | 3.1200 |
| 10.1080 | 3.1200 |
| 10.1100 | 3.1000 |
| 10.1120 | 3.1200 |
| 10.1140 | 3.0650 |
| 10.1160 | 3.0100 |
| 10.1180 | 2.7900 |
| 10.1200 | 3.0200 |
| 10.1220 | 2.8900 |
| 10.1240 | 2.7600 |
| 10.1260 | 2.9700 |
| 10.1280 | 2.9900 |
| 10.1300 | 3.0200 |
| 10.1320 | 3.0500 |
| 10.1340 | 2.9000 |
| 10.1360 | 2.8600 |
| 10.1380 | 2.8000 |
| 10.1400 | 2.7400 |
| 10.1420 | 2.9000 |
| 10.1440 | 2.9700 |
| 10.1460 | 3.0400 |
| 10.1480 | 3.0000 |
| 10.1500 | 3.0400 |
| 10.1520 | 2.9900 |
| 10.1540 | 2.9400 |
| 10.1560 | 2.9800 |
| 10.1580 | 2.8100 |
| 10.1600 | 2.8700 |
| 10.1620 | 2.9300 |
| 10.1640 | 2.7900 |
| 10.1660 | 2.8000 |
| 10.1680 | 2.8400 |
| 10.1700 | 2.8800 |
| 10.1720 | 2.8900 |
| 10.1740 | 2.9300 |
| 10.1760 | 2.9700 |
| 10.1780 | 2.8800 |
| 10.1800 | 2.9400 |
| 10.1820 | 3.0850 |
| 10.1840 | 3.2300 |
| 10.1860 | 2.7600 |
| 10.1880 | 2.8450 |
| 10.1900 | 2.9300 |
| 10.1920 | 2.7100 |
| 10.1940 | 2.8400 |
| 10.1960 | 2.7800 |
| 10.1980 | 2.7200 |

| | |
|---|---|
| 10.2000 | 2.8300 |
| 10.2020 | 3.1200 |
| 10.2040 | 3.0900 |
| 10.2060 | 3.0600 |
| 10.2080 | 2.9100 |
| 10.2100 | 2.9550 |
| 10.2120 | 3.0000 |
| 10.2140 | 2.9200 |
| 10.2160 | 2.9470 |
| 10.2180 | 2.9730 |
| 10.2200 | 3.0000 |
| 10.2220 | 3.0100 |
| 10.2240 | 2.8500 |
| 10.2260 | 2.9250 |
| 10.2280 | 3.0000 |
| 10.2300 | 2.9200 |
| 10.2320 | 2.9800 |
| 10.2340 | 2.9500 |
| 10.2360 | 2.9200 |
| 10.2380 | 2.8800 |
| 10.2400 | 2.8700 |
| 10.2420 | 2.8600 |
| 10.2440 | 2.8800 |
| 10.2460 | 2.8000 |
| 10.2480 | 2.8400 |
| 10.2500 | 2.8800 |
| 10.2520 | 2.9300 |
| 10.2540 | 2.9250 |
| 10.2560 | 2.9200 |
| 10.2580 | 2.9500 |
| 10.2600 | 3.1000 |
| 10.2620 | 3.1250 |
| 10.2640 | 3.1500 |
| 10.2660 | 3.1100 |
| 10.2680 | 3.1100 |
| 10.2700 | 3.1950 |
| 10.2720 | 3.2800 |
| 10.2740 | 3.1800 |
| 10.2760 | 3.1300 |
| 10.2780 | 3.1000 |
| 10.2800 | 3.0700 |
| 10.2820 | 2.9300 |
| 10.2840 | 2.8000 |
| 10.2860 | 2.8300 |
| 10.2880 | 2.8300 |
| 10.2900 | 2.8300 |
| 10.2920 | 2.9300 |
| 10.2940 | 2.9700 |
| 10.2960 | 2.7800 |
| 10.2980 | 2.8650 |
| 10.3000 | 2.9500 |
